# A Topological Switch Enables Misfolding of the Cystic Fibrosis Transmembrane Conductance Regulator

**DOI:** 10.1101/2020.07.09.195099

**Authors:** Daniel Scholl, Maud Sigoillot, Marie Overtus, Rafael Colomer Martinez, Chloé Martens, Yiting Wang, Els Pardon, Toon Laeremans, Abel Garcia-Pino, Jan Steyaert, David N. Sheppard, Jelle Hendrix, Cédric Govaerts

## Abstract

Cystic Fibrosis (CF) is a common lethal genetic disorder caused by mutations in the cystic fibrosis transmembrane conductance regulator (CFTR) anion channel. Misfolding and degradation of CFTR are the hallmarks of the predominant mutation, F508del, located in the first nucleotide binding domain (NBD1). While the mutation is known to affect the thermal stability of NBD1 and assembly of CFTR domains, the molecular events that lead to misfolding of F508del-CFTR remain elusive. Here, we demonstrate that NBD1 of CFTR can adopt an alternative conformation that departs from the canonical NBD fold previously observed for CFTR and other ATP-binding cassette (ABC) transporter proteins. Crystallography studies reveal that this conformation involves a topological reorganization of the β-subdomain of NBD1. This alternative state is adopted by wild-type CFTR in cells and enhances channel activity. Single-molecule fluorescence resonance energy transfer microscopy shows that the equilibrium between the conformations is regulated by ATP binding. Under destabilizing conditions, however, this conformational flexibility enables unfolding of the β-subdomain. Our data indicate that in wild-type CFTR switching to this topologically-swapped conformation of NBD1 regulates channel function, but, in the presence of the F508del mutation, it allows domain misfolding and subsequent protein degradation. Our work provides a framework to design conformation-specific therapeutics to prevent noxious transitions.

## Introduction

Cystic fibrosis (CF) is caused by mutations in the cystic fibrosis transmembrane conductance regulator (CFTR) anion channel that lead to impaired epithelial ion transport with dramatic consequences in multiple organs, particularly the lungs (Ratjen et al., 2015; Riordan et al., 1989). In most patients, CF is caused by the deletion of the phenylalanine residue at position 508 (F508del) in the first nucleotide-binding domain (NBD1), leading to absence of the channel from the plasma membrane (Cheng et al., 1990; Lukacs and Verkman, 2012). The mutation F508del compromises CFTR folding and stability through thermal destabilization of NBD1 and disruption of interdomain assembly (He et al., 2015; Protasevich et al., 2010; Serohijos et al., 2008; Wang et al., 2010). The mutant channel is recognized as misfolded at physiological temperatures, and is degraded by the cellular quality control system (Okiyoneda et al., 2010; Sharma et al., 2001). *In vitro* studies demonstrate that F508del has little effect on the NBD1 structure per se, but alters stability and dynamics of the domain (Lewis et al., 2005, 2010; Protasevich et al., 2010). Yet, the molecular events that lead to NBD1 misfolding remain poorly understood. Molecular insight can be gained from stabilizing mutations that compensate for the effects of F508del, leading to recovery of membrane expression and channel function (Aleksandrov et al., 2012; DeCarvalho et al., 2002; He et al., 2015; Lewis et al., 2005, 2010; Liu et al., 2012; Pissarra et al., 2008; Rabeh et al., 2012; Roxo-Rosa et al., 2006; Yang et al., 2018). For example, replacing residues S492, A534 and I539 in human CFTR by their avian counterparts largely suppresses the deleterious effects of F508del (Aleksandrov et al., 2012). Another striking example where a sequence change rescues the effect of the mutation involves the regulatory insertion (RI), a 32-residue long segment found in NBD1 of all CFTR orthologues, yet absent in related ATP-binding cassette (ABC) transporters. The role of the RI has remained obscure. It is not resolved in crystal structures of NBD1 (Atwell et al., 2010; Lewis et al., 2004, 2005) nor cryo-EM structures (Liu et al., 2017; Zhang and Chen, 2016; Zhang et al., 2018) of CFTR and has been described as intrinsically disordered (Lewis et al., 2010). Remarkably, removal of the RI increases the stability of NBD1 and, in the context of F508del-CFTR, largely recovers maturation, cell surface expression, activity and interdomain assembly of the mutant channel (Aleksandrov et al., 2010). Binding of ATP to NBD1, which is required for correct conformational maturation of CFTR (Lukacs et al., 1994), counterbalances destabilization by F508del to some degree (Gong et al., 2016; Wang et al., 2010). While the Walker B motif in NBD1 lacks a catalytic base (Lewis et al., 2004), making this ATP-binding site non-hydrolytic (Aleksandrov et al., 2002), ATP binding is proposed to drive opening of phosphorylated CFTR channels by promoting the formation of a NBD1:NBD2 dimer (Vergani et al., 2005). We set out to examine how factors such as ATP, stabilizing mutations or removal of the RI impact the conformational landscape of NBD1 to understand the molecular origins of CF and inform therapeutic development.

Using single molecule Förster resonance energy transfer (smFRET) and nanobody-enabled structural studies, we investigated the role of the RI in CFTR function and stability. We report that the RI enables CFTR to adopt a previously unknown conformation that modulates channel activity but also makes NBD1 vulnerable to unfolding. X-ray crystallography demonstrates that this conformation exhibits significant structural rearrangements compared to the ‘canonical’ state of NBD1 originally described. Using smFRET and hydrogen deuterium exchange coupled to mass spectrometry, we characterized the conformational equilibrium between the two states of NBD1. We show that the non-canonical conformation of NBD1 confers enhanced channel activity on wild-type CFTR. We identify the molecular determinants of the equilibrium and show that the conformational transitions also favour local unfolding of NBD1 in the presence of the F508del mutation, providing a molecular mechanism for the pathogenesis of CF.

## Results

### The Regulatory Insertion Adopts Distinct Conformational States

To characterize the conformational landscape of the RI, we used smFRET to monitor the distance between individual donor and acceptor dyes covalently attached to engineered cysteines in purified NBD1 (Fig. S1).

To overcome the poor expression and stability of NBD1 from wild-type human CFTR, we used the 2PT-NBD1 variant, which contains three stabilizing mutations found in avian CFTR (S492P, A534P and I539T) (Aleksandrov et al., 2012). Guided by previous studies of functional full-length cysteine-less CFTR variant (Cui et al., 2006; Mense et al., 2006), we replaced endogenous cysteines in 2PT-NBD1. Subsequently, for each FRET reporter, we introduced a pair of cysteines at desired locations and labelled the purified variants with ATTO488 and Alexa647 as donor and acceptor fluorophores, respectively.

To monitor movements of the RI, we chose to label position 426 located within the RI (residues 405-436, red dashed line in Fig. 1A) and residue 519 positioned within a structured α-helix. The FRET efficiency between this 426-519 reporter pair showed a broad distribution of FRET states with a major population centred around *E_FRET_* = 0.4 and two minor populations at *E_FRET_* = 0.7 and *E_FRET_* = 0.9 when measured at room temperature (22 °C) in a buffer containing 2 mM ATP (Fig. 1B). To assess whether conformational transitions occur on the submillisecond timescale, we generated two-dimensional (2D) plots of donor lifetime versus FRET efficiency. The relationship between donor lifetime and FRET efficiency depends on the kinetics of the conformational changes relative to the observation time. Specifically, if the signals fall on the predicted *static FRET line* (see Methods), there is no conformational exchange during the passage of the protein through the confocal volume of the smFRET microscope (~1–5 ms), indicating that the protein exists in one conformation at a time. By contrast, if the signals fall above the static FRET line structural changes occur, indicating dynamic behaviour on the submillisecond timescale. For the 426-519 reporter pair, we observed that the population at *E_FRET_* = 0.4 falls on the static FRET line and thus is long-lived while the populations at 0.7 and 0.9 are off the static FRET line and appear to be dynamic (Fig. 1D). To investigate whether ATP binding modulates the conformational behaviour of the RI we also performed smFRET in the absence of ATP and observed a dramatic change in the conformational profile as the entire population was shifted to *E_FRET_* values centred around 0.7 and 0.9 (Fig. 1C) and located off the static FRET line (Fig. 1E).

**Fig. 1.**
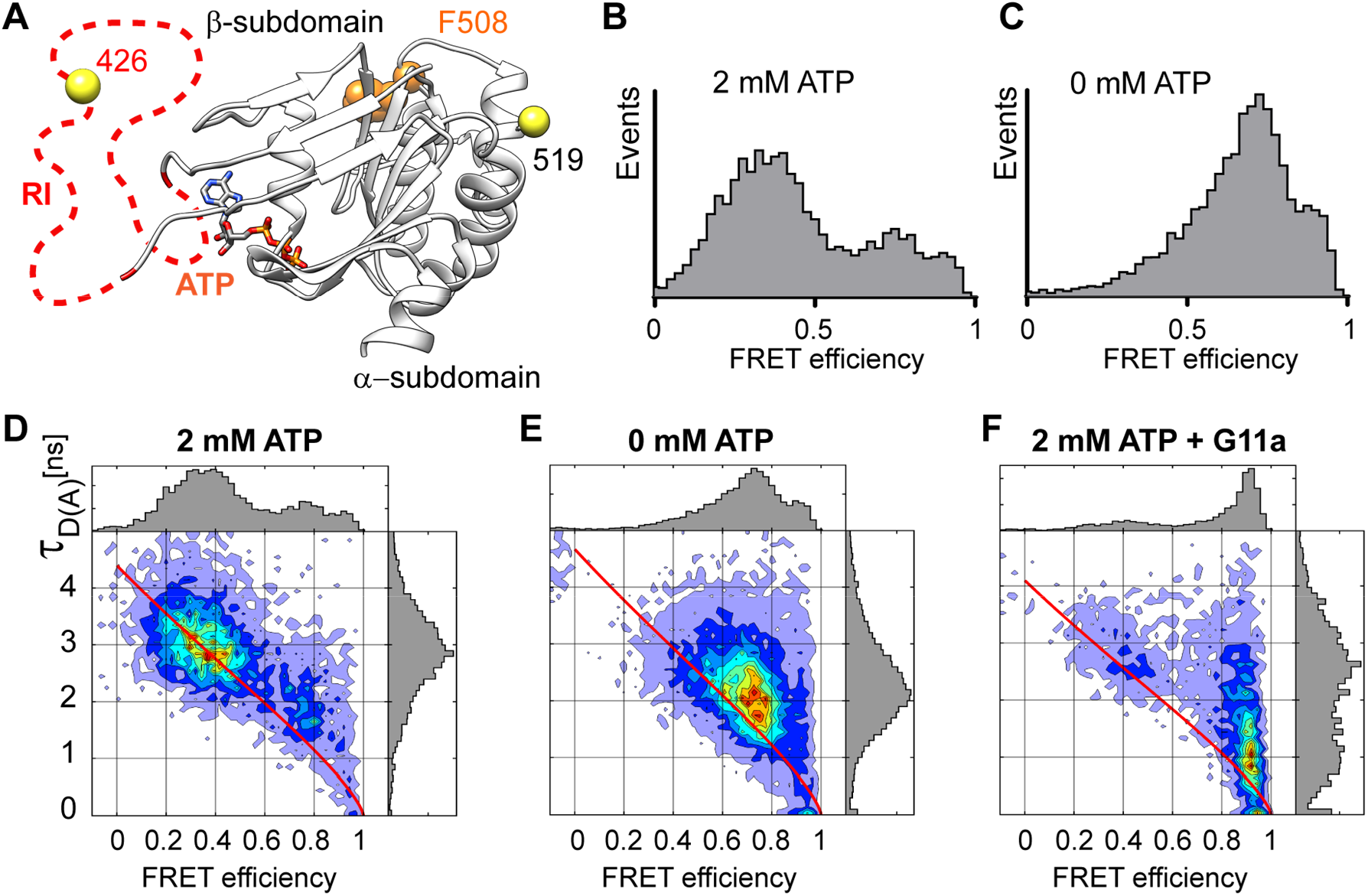
Single molecule FRET analysis for the 426-519 reporter pair of NBD1. **A,** Location of the smFRET reporters in the canonical conformation of NBD1 (based on the published structure of human NBD1, PDB: 2BBO). The positions of the labelled cysteines are shown as yellow spheres. The regulatory insertion (RI), which is not resolved in the structure, is depicted as a red dashed line. ATP and F508, located in the α-subdomain, are shown. **B-C,** FRET efficiency histograms of the 426-519 reporter in the presence and absence of 2 mM ATP and 3 mM MgCl_2_. **D-F,** Two-dimensional plots of donor fluorescence lifetime in presence of acceptor (τ_D (A)_) versus FRET efficiency of the 426-519 reporter pair for the indicated conditions. For panel **F**, in the presence of 5 μM of nanobody G11a.

To characterize the structural changes between the low and high FRET states, we screened a library of nanobodies directed against 2PT-NBD1 (Sigoillot et al., 2019) to identify RI-conformation specific binders. We identified a nanobody termed G11a which, upon binding to 2PT-NBD1, completely restricted the conformation of the RI to the high FRET state at *E_FRET_* = 0.9 (Figs. 1F). This state also existed in the absence of the nanobody (Figs. 1D-E). Thus, the nanobody did not induce new conformations of 2PT-NBD1, but rather stabilized a protein state which pre-existed in the conformational ensemble of the antigen as it was presented to the immune system of the llama used to generate the nanobodies.

To summarize, smFRET measurements demonstrated that the conformational space of the RI encompasses at least three preferred states whose relative populations are regulated by ATP binding. These can be also visualized using 2D histograms of FRET efficiency versus acceptor lifetime (Fig. S2).

### Binding of Nanobody G11a Stabilizes an Alternative Conformation of NBD1

We characterized the biophysical and biochemical effects of nanobody G11a and observed that it stabilizes 2PT-NBD1 upon binding, increasing its melting temperature by 10 °C in differential scanning fluorescence (DSF) experiments (Fig. S3A). By contrast, G11a binding was entirely lost with the ΔRI-NBD1 variant where residues 405–436 have been removed, suggesting that G11a interacts directly with part of the RI or binds a conformation adopted only in presence of the RI (Fig. S3B). We determined the high-resolution crystal structure of the 2PT-NBD1:G11a complex (Fig. 2A) at 2.7 Å (see Table S2) and found that the binding epitope includes several residues of the RI (residues 423–436, red), as well as part of strand S4 (residues 476–487, blue). The conformation of NBD1 as captured by G11a is characterized by extensive structural changes in the β-subdomain resulting in an alternative topology compared to the previously published crystal structures of NBD1 of CFTR (Lewis et al., 2004, 2005) and of NBDs from other ABC proteins (Fig. 2B-C).

**Fig. 2.**
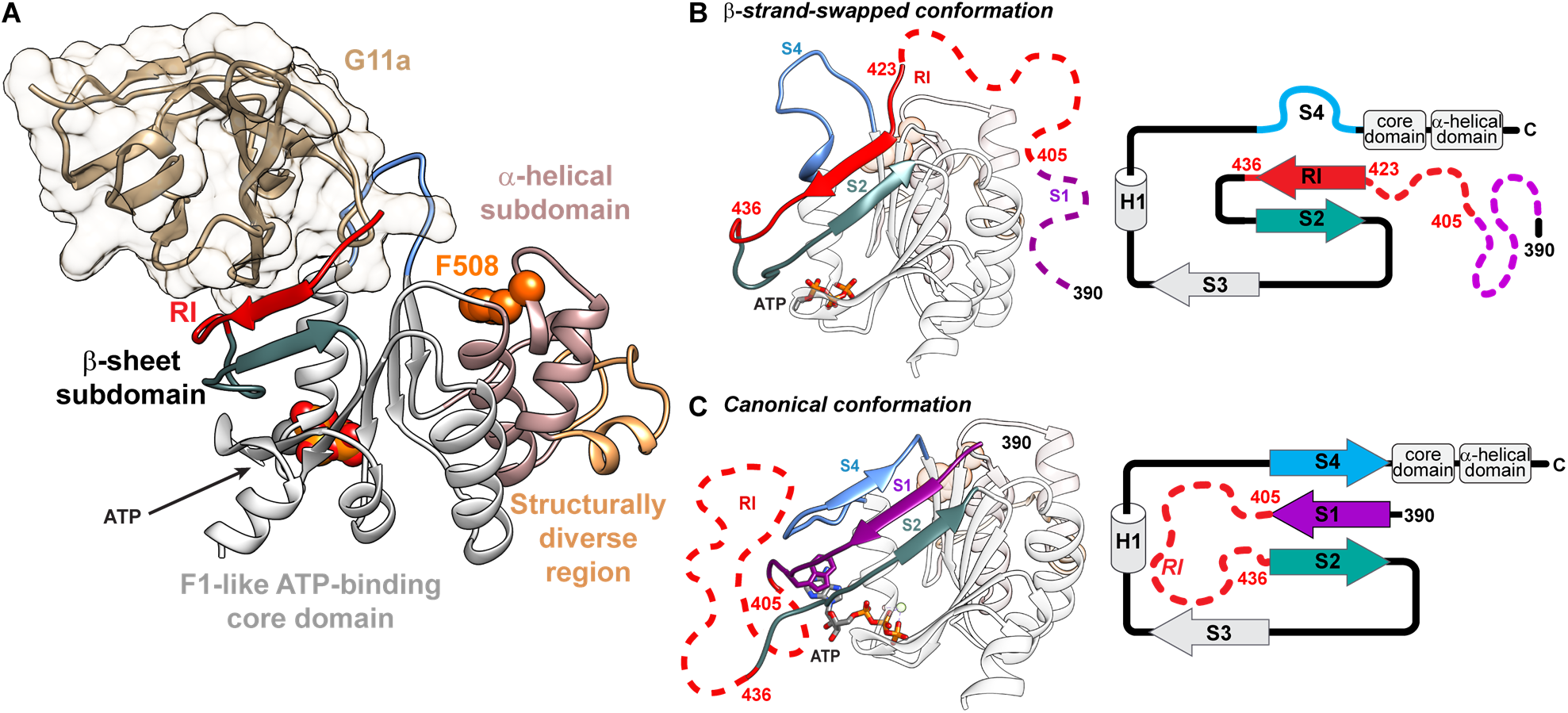
The G11a-bound structure reveals a new conformation of NBD1 with a different topology. **A,** High-resolution structure of 2PT-NBD1 in complex with nanobody G11a. No density is observed for residues 389–422. **B,** and **c,** comparative representations highlighting the topological changes in the β-subdomain observed between the G11a-bound structure (**B**, the nanobody is omitted for clarity) and the canonical conformation as previously published (**c**, here PDB: 2BBO). Unresolved segments are illustrated as random coils with dashed lines in red for the RI (404–436) and in magenta for the N-terminus (389–403). The segment including the S4 strand (473–487) is coloured in blue. A topological representation of the β-subdomain is shown to the right of each conformation. In the new conformation, residues from the RI adopt a β-sheet structure and replace the S1 segment which becomes unstructured. We therefore refer to it as *β-strand-swapped*.

All structures of human NBD1 published so far adopt a similar fold (Atwell et al., 2010; Lewis et al., 2004, 2005). In this canonical conformation (Fig. 2C), the β-subdomain is made up of three β-strands: the N-terminal S1 strand (389–404, purple), the S2 strand (437–448, turquoise), and the S4 strand (473–487, blue). The RI (405–436, red dashed line) is unstructured and connects S1 and S2. In the G11a-bound structure, we observed an alternative topology (Fig. 2B) of this β-sheet where only the S2 strand is maintained in its original position. The S4 strand (blue) has unfolded to adopt a loop structure which no longer interacts with other secondary structures of NBD1. Part of the RI has become structured and interacts with the S2 strand in an antiparallel β-sheet (red) thus replacing the original S1 strand. As a consequence, the entire 389–422 segment including the canonical S1 strand and the remainder of the RI (schematized as purple and red dashed lines in Fig. 2) is disordered in the crystal structure. As the S1 strand is replaced by a structured segment of the RI, we refer to this conformation as *β-strand-swapped* (β-SS).

Although this complex was crystallized in the presence of 2 mM ATP, we could only resolve the triphosphate of ATP with no clear density observed for the base. This observation is likely related to the fact that W401, which coordinates the adenosine base in the canonical conformation (Fig. 2C, purple), is disordered in the β-strand-swapped conformation.

In the β-SS conformation, residues 426 and 519, serving as anchor points for the FRET dyes, are separated by only 26.5 Å. Simulation of all accessible dye positions using FRET-restrained positioning and screening (FPS) software (Kalinin et al., 2012) predicts an *E_FRET_* value of 0.89 (Fig. S4A and Table S3), which agrees remarkably well with the observed value of 0.9 (Fig. 1). We conclude that the high FRET state identified by smFRET corresponds to this G11a-bound β-SS conformation characterized by crystallography.

### The β-SS Conformation is Sampled by Wild-Type CFTR and Compatible with Channel Function

The extensive and unexpected structural rearrangements observed in the β-SS conformation prompted us to question its biological relevance. As G11a binding is not compatible with the canonical conformation of NBD1, we first established whether the nanobody binds a physiologically relevant form of CFTR, an indication that the β-SS state exists naturally. To this end, we performed flow cytometry experiments on permeabilized HEK293 cells stably expressing wild-type human CFTR incubated with either nanobody G11a, nanobody T2a (a nanobody that binds a conformationally invariant epitope on NBD1 (Sigoillot et al., 2019)) or a negative control nanobody (that does not bind NBD1). Although signal intensity was reduced compared to T2a, we observed robust labelling by nanobody G11a compared to the negative control (Fig. 3A). This result indicates that the β-SS conformation can be adopted by wild-type CFTR expressed in cells where it likely represents a subpopulation of the conformational ensemble. We also performed pull-down experiments with G11a on detergent solubilized membranes expressing wild-type CFTR. SDS-PAGE and Western blotting analysis demonstrated that G11a binds both immature and mature, fully glycosylated CFTR (Fig. 3B).

**Fig. 3.**
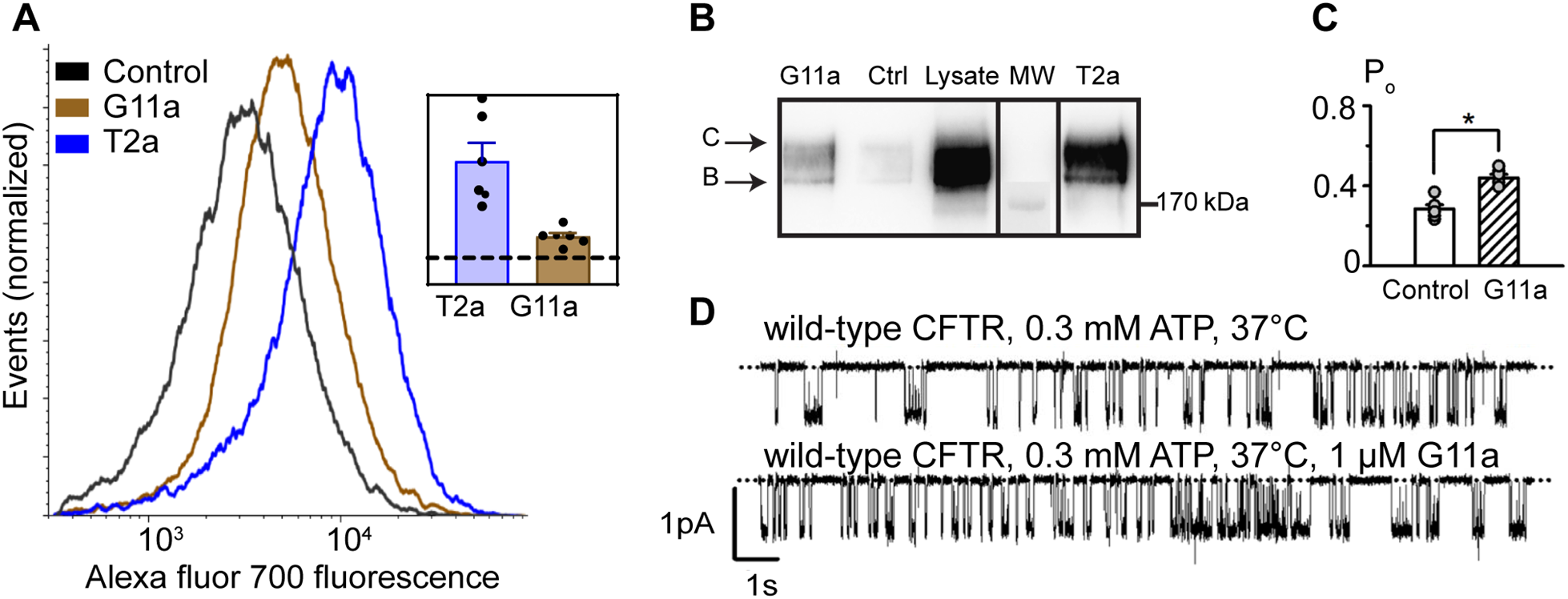
The G11a binds to full-length CFTR and increases channel activity. **A,** Flow cytometry analysis of CFTR recognition by nanobodies in CFTR overexpressing HEK293 cells. Data were normalized to the number of events acquired in each condition. Graph depicts one representative of six independent experiments. The inset shows the average median fluorescence (fold against negative control). **B,** Immunoblot of CFTR from solubilized HEK293 cells pulled down with TwinStrep-tagged nanobodies. Eluted nanobody-CFTR complexes were separated by SDS-PAGE and presence of CFTR was detected with a mouse anti-CFTR monoclonal antibody (596, recognizing residues 1204–1211 in NBD2 (Cui et al., 2007)) after immunoblotting. Arrows indicate the mature (band C) and immature (band B) forms of CFTR. **C,** Open probabilities of wild-type CFTR in the absence and presence of G11a (1 μM). Symbols represent individual values and columns are means ± SEM (n = 6); *, P < 0.05 vs control. **D,** Representative recordings of a wild-type CFTR Cl^−^ channel in an excised inside-out membrane patch from a C127 cell heterologously expressing wild-type human CFTR. The recordings were acquired at 37 °C in the presence of ATP (0.3 mM) and PKA (75 nM) in the intracellular solution. After the channel was fully activated, G11a (1 μM) was directly added to the intracellular solution bathing the membrane patch. Dotted lines indicate where the channel is closed and downward deflections correspond to channel openings.

To investigate whether the β-SS conformation of NBD1 affects the function of wild-type CFTR, we studied the single-channel activity of wild-type CFTR. Figure 3D and Figure S5 show representative recordings of an individual wild-type CFTR Cl^−^ channel in the absence and presence of saturating concentrations of G11a at 37 °C. Nanobody G11a had no effect on current flow through CFTR, but altered its gating pattern by increasing the frequency, but not the duration of channel openings, leading to a 50% increase in open probability (P_o_; a measure of the average fraction of time that a channel is open) (Fig. 3C). Analysis of prolonged recordings revealed that, in the presence of G11a, CFTR switched between two patterns of channel gating, one similar to wild-type and a second characterized by an elevated P_o_ (Fig. S5D-G). These data suggest that the alternative conformation might represent a hyperactive channel state.

We conclude that nanobody G11a exclusively stabilizes a non-canonical conformation of the RI and X-ray crystallography shows that this conformation is associated with extensive structural changes in the β-subdomain of NBD1. This alternative, *β-strand-swapped* conformation is adopted by wild-type CFTR and constitutes a functional state of the channel.

### The Conformational Equilibrium of NBD1 is Regulated by ATP

To characterize the conformational equilibrium of NBD1, we performed smFRET on a second pair of distance reporters. To follow the motions of the S4 strand, we labelled residues 479 and 519 which are separated by 41 Å in the canonical conformation (C_β_-C_β_ distance), versus 35 Å in the β-strand-swapped conformation (Fig. 4A-B). Simulations of labelling these sites yielded predicted *E_FRET_* values of 0.47 and 0.64, respectively (see Fig. S4B and Table S3).

**Fig. 4.**
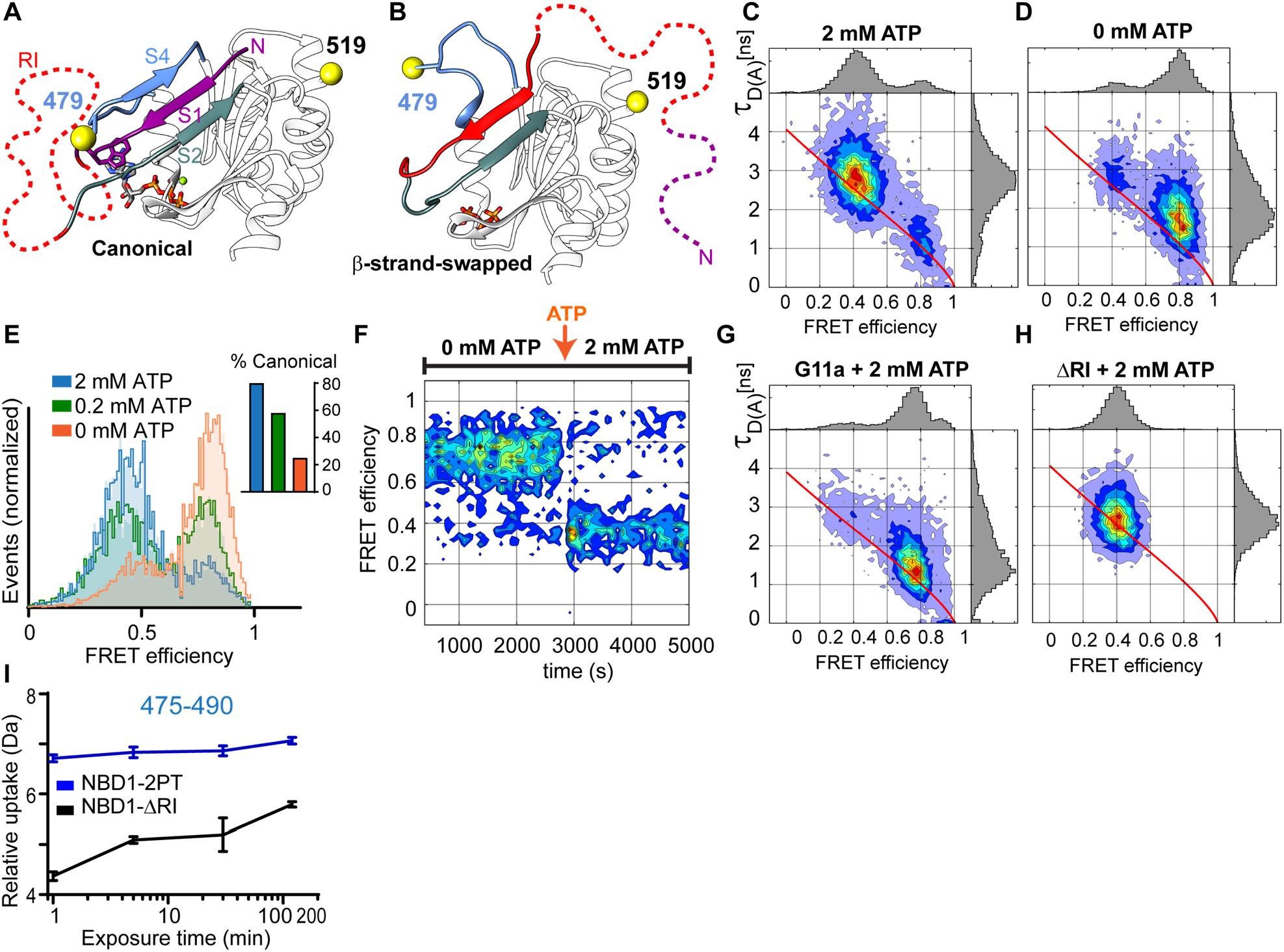
Single molecule FRET analysis for the 479-519 reporter pair of NBD1. **A-B,** Positions of the labelled cysteines 479 and 519 for the canonical and β-SS conformations. **C-D,** Two-dimensional plots of donor fluorescence lifetime (τ_D (A)_) versus FRET efficiency for the indicated constructs and conditions. In the absence of conformational dynamics, the data clouds fall on the static FRET line (red continuous line). **E,** Overlay of FRET efficiency histograms of 2PT-479-519 for the indicated ATP concentrations. Raw data (filled) were fitted using the PDAFit software (thick lines, see Methods). Inset: populations of the canonical conformation as calculated by PDAFit. **F,** Reversibility of the conformational change. **G-H,** Two-dimensional plot of 2PT-479-519 in presence of G11a (**G**) and of the ΔRI-2PT-NBD1 variant with 2 mM ATP (**H**). **I,** Deuterium uptake curves of the 475–490 peptide identified in mass spectral analyses of NBD1-2PT (blue) and ΔRI-NBD1 (black). Error bars represent the standard deviation of technical triplicates. Data shown are representative of three biological replicates.

When we measured the smFRET signal of this pair (Fig. 4C), we observed two distinct states with a major population centred on *E_FRET_* = 0.4 in agreement with predictions for the canonical conformation and a smaller population centred on 0.8 (larger than the predicted value deduced from the crystal structure of the 2PT-NBD1:G11a complex). This high FRET population increased sharply in the absence of ATP (Fig. 4D), confirming that it corresponds to a non-canonical conformation. Furthermore, we observed that the data in the absence of ATP lay off the static FRET, implying dynamic behaviour on the sub-millisecond timescale. Using photon distribution analysis (PDA), we quantified the relative populations detected by FRET using 2 mM ATP, 0.2 mM ATP and 0 mM ATP in the buffer (Fig. 4E). We observed that, at 2 mM ATP, about 80% of the distribution sampled the canonical state and 20% the putative β-SS conformation. When we lowered the ATP concentration, the population of the canonical state was decreased to 58% and 25% for 0.2 mM and 0 mM ATP, respectively, demonstrating that modulation by ATP is concentration dependent.

We then performed a kinetic experiment where, by using 0 mM ATP, the non-canonical state was initially imposed, as evidenced by the 479-519 FRET signal at *E_FRET_* = 0.8 (Fig. 4F). After 3000 s, we added ATP to a final concentration of 2 mM and observed an immediate and stable change in the FRET signal to *E_FRET_* = 0.4 indicating transition to the canonical state. These data reveal that the non-canonical conformation is not an irreversible misfolded end point of the conformation landscape, but rather an entirely reversible state.

Subsequently, we tested the effect of G11a binding on the 479-519 reporter pair using 2 mM ATP and detected a shift of the population to *E_FRET_* = 0.8 (Fig. 4G). We deduced that the high FRET state observed in the absence of the nanobody matches the opening of the S4 strand in the β-SS state captured with the G11a-NBD1 crystal. However, analysis of the FRET vs. donor lifetime 2D plot showed that this population lies on the static FRET line (Fig. 4G), indicating that binding of G11a stabilizes the opening of the S4 and restricts its dynamics.

### The RI Confers Structural Plasticity on the β-subdomain

We performed the smFRET experiments with a ΔRI-2PT variant, where the whole regulatory insertion has been removed (405-436, red in Fig. 4A-B), and observed only one population with *E_FRET_* = 0.4, matching the expected value for the canonical state (Fig. 4H).

To confirm that the RI increases plasticity of the S4 strand, we performed hydrogen deuterium exchange coupled to mass spectrometry (HDX-MS), which identifies deuterium uptake of labile protons on backbone amides to report solvent accessibility and conformational flexibility. In the presence of the RI, we observed a rapid and marked exchange of backbone amide hydrogens of the peptide 475–490, containing the S4 strand residues, indicative of increased structural flexibility (2PT-NBD1, blue; Fig. 4I). By contrast, when we performed these measurements with a ΔRI-NBD1 variant, the uptake remained limited, even after 2 h of incubation, thus confirming that RI enables flexibility of the S4 strand (ΔRI-NBD1, black; Fig. 4I).

In the β-SS conformation the N-terminal S1 strand is disordered and no longer interacts with the S2 and S4 strands of the β-subdomain (Fig. 2B). To probe the conformational equilibrium of the S1 strand, we used the 390-519 reporter pair and performed the same experiments as above (Fig. S6). We observed two FRET states that coexist and which can reliably be attributed to the canonical and β-SS conformations by using either a ΔRI variant or the G11a nanobody, respectively. As observed with the 479-519 reporter pair, the equilibrium is regulated by the presence of ATP and is reversible. HDX-MS also demonstrated that the dynamics of this region is directly coupled to the presence of the RI as we observed very limited exchange in the 392–399 peptide in ΔRI-NBD1 compared to 2PT-NBD1 (Fig. S6I). These findings are in agreement with previous HDX studies by Premchandar et al. who identified the RI, the N-terminus and the S4 strand as fast-exchanging regions (Premchandar et al., 2017).

We conclude that two conformations of the S4 and S1 segments can be distinguished by smFRET. One state corresponds to the canonical conformation where both S1 and S4 segments are engaged in the canonical β-sheet. The other state is promoted by depletion of ATP and matches predictions for the β-SS conformation observed by crystallography with an unstructured S4 and a disordered S1 segment.

### F508del Modulates the Conformational Equilibrium of NBD1

We then investigated the effects of the F508del mutation and other destabilizing factors on the conformational equilibrium of NBD1. Using the 479-519 reporter pair as a proxy and keeping a saturating concentration of ATP in the buffer, we observed that removing F508 did not affect the equilibrium at 22 °C (Fig. 5A and Table S4). However, it is known that the effects of F508del are strongly temperature-dependent as demonstrated by robust recovery of CFTR expression and function when cells expressing F508del-CFTR are grown below 30 °C (Denning et al., 1992). To test whether increasing the temperature altered the relative equilibrium populations, we performed smFRET measurements at 37 °C and observed a decrease in the canonical state of 2PT-479-519 when compared to the same construct at 22 °C (Fig. 5A), indicating that the conformational equilibrium is temperature dependent.

**Fig. 5.**
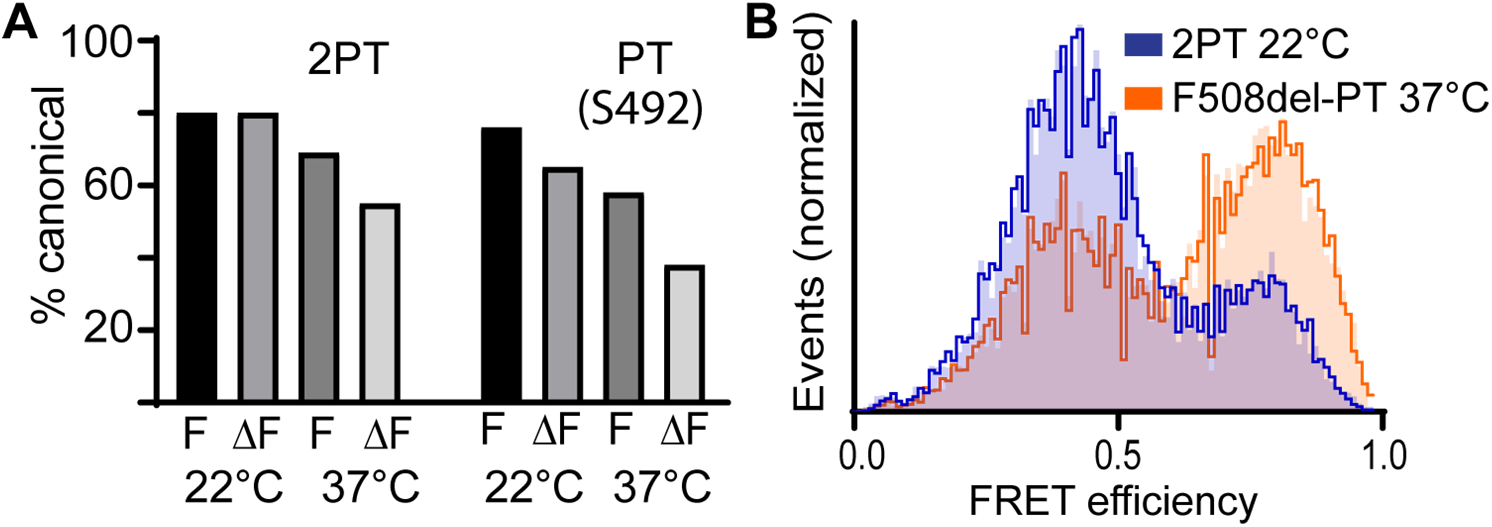
The effects of F508del, temperature and S492P on the population of the canonical state. **A,** Histograms of the population of the canonical states of the 479-519 reporter pair as calculated by PDAFit. Values are shown for 2PT-NBD1 and PT-NBD1 (with S492) variants at 22 °C and 37 °C in the presence (‘F’) or absence (‘ΔF’) of F508. **B,** Overlay of FRET efficiency histograms of 2PT-479-519 at 22 °C and F508del-PT-479-519 at 37 °C.

When we probed the conformational equilibrium at 37 °C, we observed that removal of F508del led to a marked decrease in the population of the canonical state (Fig. 5A), indicating that the CF-causing mutation modulates the equilibrium in favour of a non-canonical state at physiological temperatures.

The 2PT-NBD1 construct used here contains 3 stabilizing substitutions: S492P, A534P and I539T which reduce the impact of F508del on the maturation of the full-length protein (Aleksandrov et al., 2012). In human NBD1, Ser492 is a critical node in the allosteric network propagating thermal fluctuations between the β-subdomain and the F508 loop region (Proctor et al., 2015). To understand better the consequences of deleting F508, we analysed the impact of F508del on the PT-NBD1 construct (carrying only two stabilizing mutations: A534P and I539T), bearing S492 as in wild-type human CFTR. Using the 479-519 pair reporter, we observed a modest decrease in the population of the canonical state from 80% (2PT-NBD1) to 76% (PT-NBD1) at room temperature but a stronger decrease at 37 °C, from 69% to 58% (Fig. 5A). Finally, when combining the destabilizing factors by measuring the equilibrium in a F508del-PT-NBD1 variant at 37 °C, we observed that less than 40% of the population remained in the canonical state, despite the presence of 2 mM ATP (Fig. 5B). Fig. S7 details how each individual parameter gradually influences the conformational equilibrium.

We conclude that conditions which reduce the effects of F508del on full-length CFTR protein (i.e. low temperature or stabilizing mutations) favour the canonical state in our smFRET experiments. By contrast, deletion of F508 decreases the canonical population, but only at permissive temperatures (i.e. 37 °C).

### Unfolding of the β-subdomain Occurs in Non-Canonical States of NBD1

We surmised that the conformational plasticity of the RI and the β-subdomain plays a key role in the molecular origin of NBD1 misfolding. As we observed that the presence of the RI allows the dissociation of the S1 and S4 strands from the canonical β-sheet, we tracked domain unfolding by monitoring conformational changes in the rest of the β-sheet, specifically strand S2. Motion of this region is relevant to pathogenesis as residue K447 located in the S2 strand (Fig. 6A) is SUMOylated in F508del-NBD1, but not in WT-NBD1 (Gong et al., 2016), suggestive of increased exposure in mutant CFTR. We monitored the conformations of the S2 strand in smFRET by labelling residues 442 and 519. Importantly, residue 447 (and 442) is not involved in the structural rearrangements that occur between the canonical and β-SS states (Fig. 2B-C). At 22 °C, the smFRET signal of the 442-519 reporter pair in the context of 2PT-NBD1 showed a major static population with *E_FRET_* centred on 0.4 (Fig. 6B) which agrees with the expected value based on either crystal structures (Table S3). In addition to this low FRET state, we observed a minor diffuse population with *E_FRET_* values above 0.5. This indicates the coexistence of conformations where residue 442 approaches residue 519, a condition that requires local unfolding of the S2 strand. When the temperature was increased to 37 °C, 2PT-NBD1 showed a marked increase in this high *E_FRET_* population (Fig. 6D). This putatively unstructured population disappears entirely when the RI is absent (Fig. 6C), suggesting that it is linked to non-canonical states.

**Fig. 6.**
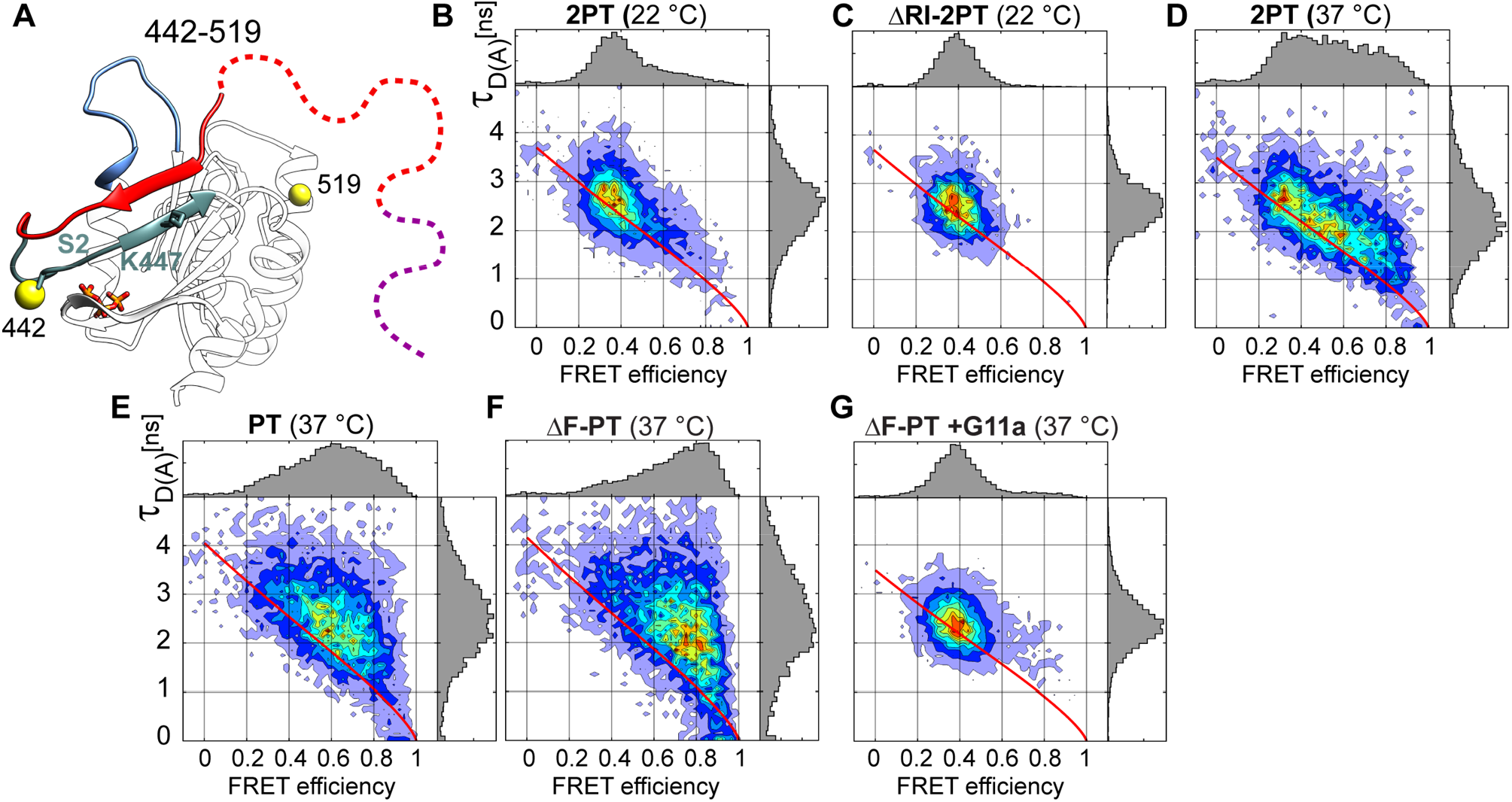
Dynamics of the S2 strand are affected by F508del. **A,** Structure of NBD1 in the β-strand swapped conformation illustrating the positions of residues 442 and 519 (yellow spheres) along with K447 in the S2 strand (turquoise). **B-G**, Two-dimensional plots of donor fluorescence lifetime (τ_D (A)_) vs FRET efficiency for the 442-519 reporter pair for the indicated constructs and conditions.

When we measured the FRET signal of the 442-519 reporter pair at 37 °C in the PT-NBD1 variant, the proportion of high FRET population increased compared to 2PT-NBD1 (Fig. 6E). Thus, the presence of S492, which decreased the population of canonical conformers (Fig. 5), also favoured local unfolding of the β-sheet.

Finally, upon deletion of F508, we observed a further increase in the high FRET population with fluorescence signals lying off the static FRET line (Fig. 6F), indicative of highly dynamic behaviour. The broad distribution of *E_FRET_* values, from 0.5 to 1, is suggestive of a disordered state. By contrast, preincubation of the F508del-PT variant with G11a completely prevented local unfolding of the β-subdomain at 37 °C as the entire population became static with *E_FRET_* = 0.4 (Fig. 6G). This matches the expected values for the β-SS conformations and agrees with the thermal stabilization of NBD1 upon binding of G11a. This suggests that binding of G11a traps NBD1 in the β-SS state and protects the β-subdomain from unfolding.

## Discussion

Our data demonstrate that CFTR alternates between the state observed by cryo-EM where NBD1 adopts the canonical conformation (Fig. 7A), and one (or more) non-canonical state (s) where NBD1 adopts a conformation which has not been observed previously (Fig. 7B). Transitions towards the latter require the RI, a unique 32-residue intrinsically disordered segment (Kanelis et al., 2010; Lewis et al., 2010). Single molecule FRET measurements showed that the conformational equilibrium of NBD1 is regulated by ATP binding, which stabilizes the canonical state. A *β-strand-swapped* state was trapped with nanobody G11a and resolved by crystallography, revealing dramatic topological changes whereby the RI rearranges into a β-strand with the S4 loop disengaging from the β-sheet, and the 389–422 segment becoming unstructured (Figs. 2B and 7B).

**Fig. 7.**
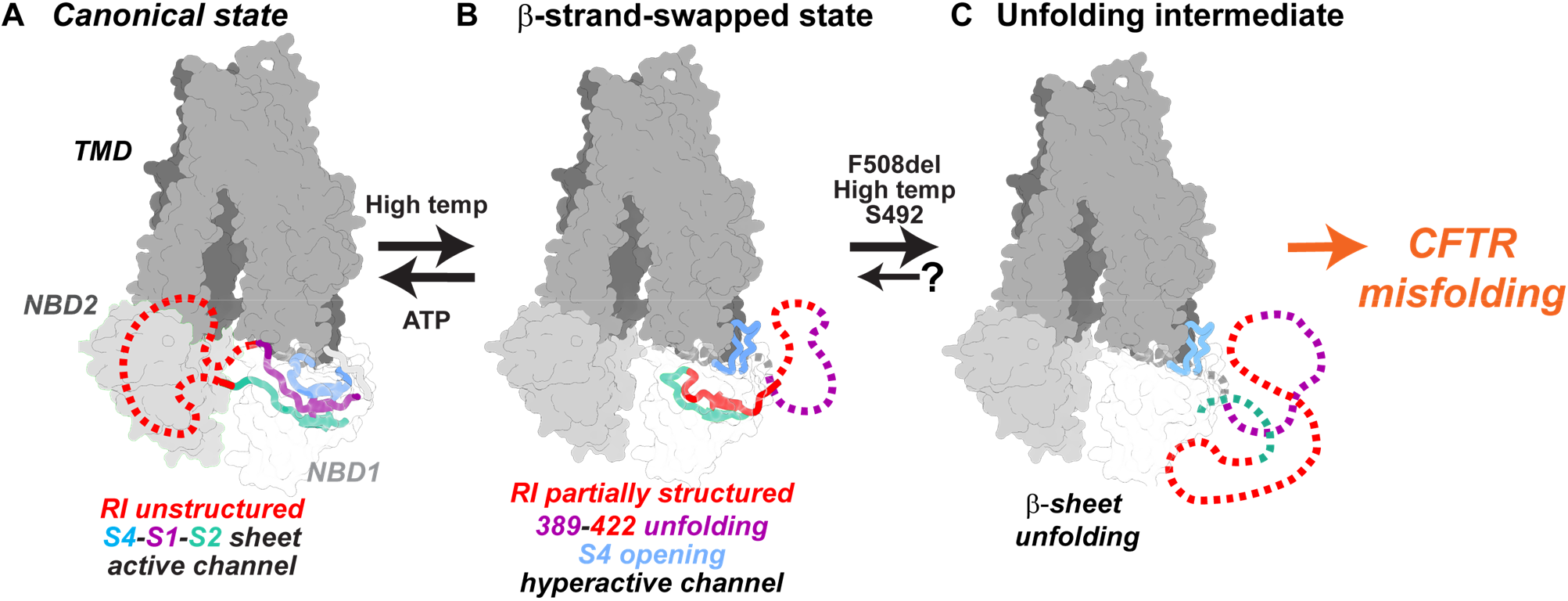
Model of an unfolding pathway through the β-strand-swapped conformation. Cartoon representations of the different conformational states of NBD1 superimposed onto the dephosphorylated and ATP-free full-length CFTR structure (PDB: 5UAK). **A,** The canonical conformation of human NBD1 (PDB: 2BBO) is mostly structured, only the RI (red) is unstructured. **B,** In the β-strand-swapped conformation, the N-terminus and the first half of the RI are unstructured (purple and red dashed, respectively). The second part of the RI is structured (red thick) and the S4 strand unfolds to become a loop (blue). **C,** In the unfolding intermediate, the RI and all strands of the β-sheet are unstructured (S1 in purple, RI in red, S2 in dark green and S4 in blue). Figures of full-length CFTR structures were created using ChimeraX (Goddard et al., 2018).

Electrophysiological measurements showed that the *β-strand-swapped* state leads to enhanced CFTR channel activity, suggesting that it might have evolved for functional reasons. Indeed, the high conservation of the RI in all CFTR orthologues and its absence in other ABC transporters endows a functional role specific to CFTR, possibly as a conformational switch (i.e. between low and high channel activities). As such, the phosphorylation site within the RI (at position S422) might serve as a regulator of the conformational switch. However, the functional gain provided by the RI seems to have come at the cost of increased vulnerability to misfolding, especially in the presence of destabilizing mutations such as F508del. Indeed, our smFRET measurements show that conditions which are known to compensate for the effects of the pathogenic mutation *in cellulo* (such as stabilizing mutations, decreased temperatures, or removal of the RI) are associated with stabilization of the canonical population of NBD1. By contrast, we observed that introducing F508del in a permissive context (37 °C, non-stabilized variant) leads to loss of the canonical conformer, which was correlated with unfolding of the β-subdomain, as measured by motion of the S2 strand (Fig. 6).

Therefore, we surmise that the protein naturally alternates between two functional states, the canonical conformation observed previously, and the β-strand-swapped conformation. In a structurally compromised context, such as destabilization by F508del, the reorganization of the β-subdomain, which characterizes the β-SS conformation might facilitate local unfolding (or prevent correct folding during biogenesis), leading to misfolding of the domain and thus eventually the entire protein.

We propose that destabilizing factors such as F508del, facilitate the transition from the β-SS state to an unfolding intermediate (Fig. 7C) where the entire β-subdomain is unfolded as measured by the motion of residue 442 (Fig. 6F). As the unfolding intermediates accumulate, the canonical population is decreased (Fig. 5) and transition to a fully misfolded state is promoted (Fig. 7C). The idea that F508del leads to unfolding in the β-subdomain is supported by previous HDX studies of NBD1 featuring only one stabilizing mutation (Premchandar et al., 2017). In these studies, Premchandar et al. showed that removal of F508 led to substantially different exchange rates only in the region around F508 itself, the N-terminus and the S2 strand (Premchandar et al., 2017).

Importantly, as this unfolding pathway requires the presence of the RI, this model explains why its removal is protective against F508del (Aleksandrov et al., 2010). Likewise, it provides an explanation for protection by S492P (Aleksandrov et al., 2012) (and possibly other revertant mutations) as the mutation allosterically limits the conformational transitions proposed above.

Previous models have proposed unfolding pathways for F508del-NBD1 in the absence of the RI (Protasevich et al., 2010), suggesting that the destabilizing effects of the deletion affect the integrity of the canonical conformation of NBD1. Our data suggest that, in a physiological context (where the RI is present), another pathway must be considered. In this pathway, F508del primarily topples CFTR from a topologically modified conformation that was not observed previously and relies on the RI.

The discovery of a new NBD1 conformation with pathophysiological implications changes our understanding of CFTR biology. It might also open conformation-specific therapeutic approaches for F508del and possibly other related CF-causing mutations. Our study sheds a new light on how structured and unstructured regions of a protein can interchange for functional reasons and might constitute a new mechanism of modulation of protein function that could extend to other biological systems.

## Methods

### Human NBD1 expression and purification

Human 2PT-NBD1 (residues 387-646 containing the mutations S492P, A534P, I539T) and ΔRI-NBD1 (residues 387-646, Δ405-436) were expressed as N-terminal, His_6_-SUMO fusion proteins in *Escherichia coli* (BL21 (DE3) pLysS cells) as previously described (Lewis et al., 2004; Protasevich et al., 2010). Cells were lysed using a French press and recombinant proteins were purified by nickel ion affinity chromatography (HisTrap HP, 1ml - GE Healthcare). The His_6_-SUMO tag was removed using Ulp1 protease (Li and Hochstrasser, 1999) followed by a second nickel ion chromatography. The samples were then passed through a gel filtration column (Superdex 200 Increase 10/300 GL - GE Healthcare) previously equilibrated in 20 mM Hepes pH 7.5, 150 mM NaCl, 10% (w/v) glycerol, 10% (w/v) ethylene glycol, 2 mM ATP, 3 mM MgCl_2_, 1 mMTris (2-carboxyethyl)phosphine (TCEP). Protein concentration was determined using Coomassie Plus (Bradford) Assay Kit (ThemoScientific).

### Nanobodies cloning, expression and purification

Nanobodies were cloned in pXAP100 vector encoding His_6_-tag and Myc-tag at the C-terminal end of the protein. This vector was also modified to encode Twin-Strep-tag (Trp-Ser-His-Pro-Gln-Phe-Glu-Lys repeated twice with an internal linker region) instead of the His_6_-tag, which was used for CFTR pull-down experiments. The expression and purification of nanobodies was performed as previously described (Pardon et al., 2014). Briefly, nanobodies were produced in *Escherichia coli (*BL21 (DE3) pLysS cells), purified from the periplasmic extract via either HisPur Ni-NTA resin (ThemoScientific) or Strep-Tactin XT Superflow resin (Iba) followed by a size exclusion chromatography (SEC) (Superdex 200 Increase 10/300 GL -GE Healthcare) into 20 mM HEPES pH 7.5, 150 mM NaCl, and 10% (w/v) glycerol.

### NBD1-ELISA assay

For dose-response assays, Nunc MaxiSorp 96-well plates (ThermoScientific), were coated with 5 μg/ml NeutrAvidin Biotin-Binding protein (ThermoScientific) overnight at 4 °C and blocked for 2 h at room temperature (RT) with 4% milk in phosphate-buffered saline (PBS). Each new reagent addition was preceded by three washes with 200 μl of NBD1 buffer (20 mM HEPES pH 7.5, 150 mM NaCl, and 10% (w/v) glycerol, 10% (w/v) ethylene glycol, 2 mM ATP, 3 mM MgCl_2_). Then, biotinylated purified NBD1 proteins at 5 μg/ml were immobilized for 45 min at RT followed by 1 h RT incubation with 100 μl various concentrations (0-100 μg/ml) of purified nanobody. Signal detection was followed using His_6_-tag specific antibody (0.5 μg/ml - Invitrogen) to detect the nanobodies and secondary antibody anti-mouse coupled to horse radish peroxidase (HRP) (0.5 μg/ml – Millipore). 50 μl of 1-Step UltraTMB-ELISA (ThermoScientific) was used as a substrate for the peroxidase and intensity of the reaction was proportional to absorbance measured at 450 nm with SynergyMx (BioTek) after addition of 50 μl H_2_SO_4_ at 1M.

### Thermal shift assay (DSF)

Solutions of NBD1 constructs (10 μM final concentration), nanobodies (30 μM final concentration) and 2.5 x concentrated SYPRO Orange Protein Stain (Molecular Probes) diluted in 20 mM HEPES pH 7.5, 150 mM NaCl, 3 mM MgCl_2_, 2 mM ATP and 10% (w/v) glycerol, 10% (w/v) ethylene glycol, were added to the wells of 96-well PCR plates (VWR) in a final volume of 25 μl. Plates were sealed with EasySeal sheets (Molecular Dimensions) and spun for 2 min at 900 x g. SYPRO orange fluorescence was monitored in CFX96 Touch Real-Time PCR Detection System (Bio-Rad) using plate type BR white and scan mode FRET from 10 to 80 °C in increments of 1 °C (Lo et al., 2004; Niesen et al., 2007).

### Crystallization trials

For 2PT-NBD1:G11a complex formation, G11a nanobody was SEC purified the day before in 20 mM Hepes pH 7.5, 150 mM NaCl, 10% (w/v) glycerol and mixed with freshly SEC purified NBD1 with 1.2 molar excess of G11a, and keeping 2 mM ATP, 3 mM MgCl_2_ and 1 mM TCEP final concentrations. Protein complex was incubated overnight on ice and then concentrated onto 30 kDa MWCO Amicon concentrator (Millipore) until protein concentration reached 11 mg/ml. Crystallization was performed in sitting drops at room temperature, adding 100 nl of protein to 100 nl of the precipitant (0.2 M CaCl_2_, 0.1 M Tris pH 8.5, 25 % (w/v) PEG 4000 and were set up immediately using Mosquito robot (Art Robbins). Crystallization plates were incubated at 20 °C. Single crystals were mounted in CryoLoops (Molecular Dimensions Ltd) and flash-frozen in liquid nitrogen.

### Crystal structure determination

Diffraction data were collected at 100 K at Proxima1 beamline of Soleil (Gif-sur-Yvette, France) synchrotron as indicated in Table S2. Data were processed using the Autoproc program (Vonrhein et al., 2011). The dataset was solved by molecular replacement using Molrep (Vagin and Teplyakov, 1997). Subsequently, several cycles of model building, using COOT (Emsley et al., 2010), combined with refinement using BUSTER 2.10.1 (Bricogne et al., 2017) were conducted. Finally, structure validation was performed with MOLPROBITY (Chen et al., 2010). Figures and structural comparisons of the 2PT-NBD1:G11a complex with the human NBD1 structure previously published (PDB: 2BBO (Lewis et al., 2010)) were prepared using Chimera (Pettersen et al., 2004).

### Flow cytometry

HEK293 cells stably overexpressing human wt-CFTR (Hildebrandt et al., 2015) were permeabilized with 0.01% n-Dodecyl-β-D-Maltopyranoside (β-DDM - Inalco) at least for 2 h on ice. In the meantime, cells were incubated with 50 μg/ml nanobodies and DAPI (2.5 μM – Invitrogen) to monitor the permeabilization state. Nanobody binding was detected using Myc-tag specific antibody (2 μg/ml – Life Technologies) and then anti-mouse-Alexa Fluor 700 (1.3 μg/ml – Invitrogen) at least for 30 min on ice. We used the T2a nanobody as positive control for CFTR binding (Sigoillot et al., 2019) and a nanobody directed against the *Lactococcus lactis* multidrug resistance protein (LmrP) as negative control. Cells were washed once between each step by centrifugation (200 x g for 5 min at 4 °C). All incubations (100 μl) and washes (1.5 ml) were performed in PBS with 6% fetal bovine serum (FBS) and 0.01% β-DDM on ice. Cells fluorescence was measured with Gallios Flow Cytometer (Beckman Coulter). Data were analyzed with Kaluza software.

### CFTR pull-down

Human wt-CFTR was extracted from HEK293 cells pellet by solubilization with 0.1% DMNG in PBS with protease inhibitors for 1 h at 4 °C. The cell debris was removed by centrifugation (16,000 x g for 30 min at 4 °C). Supernatant was diluted 10 times in PBS with protease inhibitors and incubated at least for 30 min on 10 μl Strep-Tactin XT Superflow resin (Iba) pre-loaded with approximately 200 μg nanobodies. Resin was washed and eluted with Buffer BXT (Iba). Presence of CFTR in each sample was detected by SDS-PAGE and immuno-blotting.

### SDS-PAGE and immunoblotting

Cell extracts were separated by SDS-PAGE on 7.5% polyacrylamide gel and transferred to nitrocellulose membrane (Bio-Rad) for immunodetection. After blocking for 1 h with 5% bovine serum albumin (BSA) in Tris-buffered saline added Tween-20 (TBST), CFTR was detected using monoclonal antibody 596 (Cui et al., 2007) for 1 h in blocking buffer. Blots were washed 3 times for 5 min and incubated with anti-mouse-HRP antibody (0.2 μg/ml – Millipore) for 1 h in TBST. Membranes were washed 3 times for 5 min. CFTR was visualized by chemiluminescence using Luminata Forte Western HRP Substrate (Millipore) and detected with ImageQuant 400 (GE Healthcare).

### Patch-clamp electrophysiology

#### Cells and cell culture

For patch-clamp experiments, we used mouse mammary epithelial (C127) cells stably expressing wild-type human CFTR (Marshall et al., 1994). These cells were a generous gift of C. R. O’Riordan (Sanofi Genzyme). They were cultured and used as described previously (Sheppard and Robinson, 1997).

#### Patch-clamp experiments

CFTR Cl^−^ channels were recorded in excised inside-out membrane patches using an Axopatch 200B patch-clamp amplifier and pCLAMP software (version 10.4) both from Molecular Devices (San Jose, CA, USA) as described previously (Sheppard and Robinson, 1997). The pipette (extracellular) solution contained (mM): 140 N-methyl-D-glucamine, 140 aspartic acid, 5 CaCl_2_, 2 MgSO_4_ and 10 N-tris[hydroxymethyl]methyl-2-aminoethanesulphonic acid (TES), adjusted to pH 7.3 with Tris ([Cl^−^], 10 mM). The bath (intracellular) solution contained (mM): 140 NMDG, 3 MgCl_2_, 1 CsEGTA and 10 TES, adjusted to pH 7.3 with HCl ([Cl^−^], 147 mM; free [Ca^2+^], < 10^−8^ M) and was maintained at 37 °C using a temperature-controlled microscope stage (Brook Industries, Lake Villa, IL, USA).

After excision of inside-out membrane patches, we clamped voltage at −50 mV and added the catalytic subunit of protein kinase A (PKA; 75 nM) and ATP (1 mM) to the intracellular solution within 5 minutes of membrane patch excision to activate CFTR Cl^−^ channels. To test the effects of nanobodies on the single-channel behaviour of wild-type CFTR, we reduced the ATP concentration to 0.3 mM before adding them to the intracellular solution and acquiring 5 – 10 minutes of single-channel data in the continuous presence of ATP (0.3 mM) and PKA (75 nM). Because of the difficulty of removing nanobodies from the recording chamber, specific interventions were compared with the pre-intervention control period made with the same concentration of ATP and PKA, but without nanobodies. To minimise rundown, PKA and ATP were added to all intracellular solutions.

In this study, we used excised inside-out membrane patches containing ≤ 5 active channels. To determine channel number, we used the maximum number of simultaneous channel openings observed during an experiment (Cai et al., 2006). To minimise errors, we used experimental conditions that produced robust channel activity and verified that recordings were of sufficient length to determine the correct number of channels (Venglarik et al., 1994).

Single-channel currents were acquired directly to computer hard disc after filtering at a corner frequency (*f*_c_) of 500 Hz using an eight-pole Bessel filter (model F-900C/9L8L, Frequency Devices Inc., Ottawa, IL, USA) and digitized at a sampling rate of 5 kHz using a DigiData 1440A interface (Molecular Devices) and pCLAMP software (version 10.4). To measure single-channel current amplitude (*i*), Gaussian distributions were fit to current amplitude histograms. For open probability (*P*_o_) measurements, lists of open- and closed-times were generated using a half-amplitude crossing criterion for event detection and dwell time histograms constructed as previously described (Sheppard and Robinson, 1997); transitions < 1 ms were excluded from the analysis (eight-pole Bessel filter rise time (*T*_10-90_) ~0.73 ms at *f*_c_ = 500 Hz). Histograms were fit with one- or two-component exponential functions using the maximum likelihood method. For burst analyses, we used a *t*_c_ (the time that separates interburst closures from intraburst closures) determined from closed time histograms (Cai et al., 2006). The mean interburst interval (*T*_IBI_) was calculated using the equation (Cai et al., 2006):

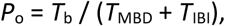

Where, *T*_b_ = (mean burst duration) × (open probability within a burst). Mean burst duration (*T*_MBD_) and open probability within a burst (*P*_o (burst)_) were determined directly from experimental data using pCLAMP software. Only membrane patches that contained a single active channel were used for burst analyses. For the purposes of illustration, single-channel recordings were filtered at 500 Hz and digitised at 5 kHz before file size compression by 5-fold data reduction.

#### Reagents

PKA purified from bovine heart was purchased from Calbiochem (Merck Chemicals Ltd., Nottingham, UK). All other chemicals were supplied by Sigma-Aldrich Ltd. (now Merck Life Science UK Ltd.) (Gillingham, UK). Stock solutions of ATP were prepared in intracellular solution directly before each experiment. Immediately before use, stock solutions were diluted to final concentrations with intracellular solution and, where necessary, the pH of the intracellular solution was readjusted to pH 7.3 to avoid pH-dependent changes in CFTR function (Chen et al., 2009). On completion of experiments, the recording chamber was thoroughly cleaned before re-use (Wang et al., 2014).

#### Statistics

Data recording and analyses were randomised, but not blinded. Results are expressed as means ± SEM of n observations, where n represents the number of individual membrane patches obtained from different cells. All data were tested for normal distribution using a Shapiro-Wilk normality test. To test for differences between two groups of data acquired within the same experiment, we used Student’s paired *t*-test. Tests were performed using SigmaPlot™ (version 13.0, Systat Software Inc., San Jose, CA, USA). Differences were considered statistically significant when *P* < 0.05. To avoid pseudo-replication, all experiments were repeated at different times.

### Hydrogen deuterium exchange mass spectrometry

#### Experiments

Hydrogen deuterium exchange mass spectrometry (HDX-MS) experiments were performed on a Synapt G2Si HDMS coupled to an Acquity UPLC M-Class system with HDX and automation (Waters Corporation, Manchester UK). Protein samples were used at a concentration of 50 μM. Isotope labelling was initiated by diluting 5 μl of each protein sample into 95 μl of buffer L (HEPES 20 mM pD 7.5, NaCl 100 mM) complemented with 2 mM ATP, 3 mM MgCl_2_ when specified. The protein was incubated for 1 min, 5 min, 30 min and 120 min to capture short, medium and long exchange times and then quenched in buffer Q (100 mM potassium phosphate, brought to pH 2.3 with formic acid) at 4 °C before being digested on-line with a Waters Enzymate BEH pepsin column at 20 °C, for ~30 s, at a pressure of ~2000 psi. The same procedure was used for undeuterated controls, with the labelling buffer being replaced by buffer E (HEPES 20 mM pH 7.5, NaCl 100 mM without 2 mM ATP and 3 mM MgCl_2_). The peptides were trapped on a Waters BEH C18 VanGuard pre-column for 3 minutes at a flow rate of 200 μl / min in buffer A (0.1% formic acid ~pH 2.5) for desalting before being applied to a Waters BEH C-18 analytical column. Peptides were eluted with a linear gradient of buffer B (8-40% gradient of 0.1% formic acid in acetonitrile) at a flow rate of 40 μl / min. All trapping and chromatography steps were performed at 1 °C to minimize back exchange. The electrospray ionization source was operated in the positive ion mode and ion mobility was enabled for the MS instrument. HDMS^E^ data were acquired with a 20 to 30 V trap collision energy ramp for high-energy acquisition of product ions. Optimized peptide identification and peptide coverage for all samples was performed from undeuterated controls (five replicates). All deuterium time points were performed in triplicate. Leucine Enkephalin (LeuEnk - Sigma) was used as a lock mass for mass accuracy correction and the MS was calibrated with sodium iodide. The on-line Enzymate pepsin column was washed between each injection with pepsin wash (1.5 M Gu-HCl, 4% MeOH, 0.8% formic acid) recommended by the manufacturer to prevent significant peptide carry-over from the pepsin column and a blank run was performed between each technical triplicate. Measurements were performed on three biological replicates.

#### Data evaluation and statistical analysis

Sequence identification was made from MS^E^ data from the undeuterated samples using ProteinLynx Global Server 2.5.1 (PLGS Waters Corp. Manchester UK). The output peptides were filtered using DynamX (v. 3.0) using the following filtering parameters: minimum intensity of 1000, minimum and maximum peptide sequence length of 5 and 25 respectively, minimum MS/MS products of 2, minimum products per amino acid of 0.28, minimum score of 5, and a maximum MH^+^ error threshold of 15 ppm. The peptides had to be identified in 3 out of 5 replicates. Additionally, all the spectra were visually examined and only those with high signal to noise ratios were used for HDX-MS analysis. The amount of relative deuterium uptake for each peptide was determined using DynamX (v. 3.0) and are not corrected for back exchange since only relative differences were used for analysis and interpretation and there was no benefit from normalizing the data (Wales et al., 2013). The relative fractional uptake (RFU) was calculated from RFU_a_ *=* [Y_a,t_/ (MaxUptake_a_ × D)], where *Y* is the deuterium uptake for peptide *a* at incubation time *t*, and *D* is the percentage of deuterium in the sample after mixing the protein with the labeling solution. 99% Confidence intervals were calculated using Deuteros (Lau et al., 2019) and were found to be of 0.7 Da or smaller for the sum of RFU over all time points.

### Modeling of dye distributions using the FPS simulation program

We used the FRET-restrained positioning and screening (FPS) software (Kalinin et al., 2012) to calculate accessible volumes and theoretical approximations of distances and FRET efficiencies between fluorophores fixed onto cysteines in key regions of NBD1. Starting from a PDB file of NBD1 (2BBO for the canonical conformation and the 2PT-NBD1:G11 complex for the β-strand swapped conformation) the program computes an accessible volume for each fluorophore based on all possible dye positions within the linker length from the attachment point excluding positions that lead to steric clashes with the protein surface. Before computing we mutated the residues to be labeled into cysteines *in silico* and attached the fluorophore to the Sulfur atom. The parameters corresponding to the ATTO488 fluorophore are 18.7 Å in length, 4.5 Å width and dye radii of 5 Å, 4.5 Å and 1.5 Å and for Alexa647: 20.3 Å in length, 4.5 Å width, and dye radii of 11 Å, 4.7 Å and 1.5 Å. We computed position 1 with ATTO and position 2 with Alexa647, using a Förster radius (R_0_) of 56.8 Å determined in (Vandenberk et al., 2018).

### Site-directed mutagenesis for the generation of single molecule PIE-FRET constructs

Guided by previous studies of functional full-length cysteine-less CFTR variants (Cui et al., 2006; Mense et al., 2006), we replaced endogenous cysteines of 2PT-NBD1. Cysteines 491 and 524 were converted to alanine and cysteines 590 and 592 to valine. Substitutions were done by the QuikChange™ site-directed mutagenesis procedure using a modified protocol for primer design (Liu and Naismith, 2008).

### NBD1 fluorophore labeling

For smFRET experiments engineered cysteines of NBD1 constructs were labeled with ATTO488 and Alexa647 maleimide dyes (ATTO-TEC GmbH AD 488-41 and ThermoFisher A20347, respectively). Affinity-purified NBD1 at 70 μM in labeling buffer (20 mM HEPES pH 7.20-7.40, 150 mM NaCl, 10% (w/v) glycerol, 10% (w/v) ethylene glycol 2 mM ATP, 3 mM MgCl_2_ and 1 mM TCEP) was incubated with a 3 times molar excess of each dye to NBD1 for a total of 3 h. Six additions of both dyes at 0.5 molar excess were administrated in 30 minute intervals to reduce preferential labeling by one dye or the other. The first two additions were performed at room temperature and the remaining ones on ice. Labeled NBD1 was spun at 4 °C and 16,000 g for 30 minutes and passed through a gel filtration column (Superdex 200 Increase 10/300 GL - GE Healthcare) previously equilibrated in 20 mM HEPES pH 7.5, 150 mM NaCl, 10% (w/V) glycerol, 10% (w/V) ethylene glycol, 2 mM ATP, 3 mM MgCl_2_.

### Single-molecule FRET data recording

The stored protein at 5-15 μM was diluted 1000-fold into the buffer (20 mM HEPES, pH 7.5, 150 mM NaCl) prior to mixing with other reagents. Then the protein was further diluted 100-fold in the final measurement sample containing also 0.1 mg/mL BSA, MgATP and/or nanobody, reaching a final concentration of 50-150 pM. The BSA-coated (1 mg/ml BSA) coverslip (Nunc Lab-Tek Chambered Coverglass, Thermo Fisher Scientific) was rinsed two times with the sample solution (30 μl), prior to depositing a 30 μl drop of the same solution. Background (needed for calculating *E* and *S* parameters, and for lifetime and PDA analysis) or scatter samples (needed for lifetime analysis) were prepared similarly but without protein.

Per sample, 1-3 hour datasets (10 minutes for background samples) were recorded at 25 °C on a homebuilt multiparameter fluorescence detection microscope with pulsed interleaved excitation (MFD-PIE) as established (Talavera et al., 2018), with minor modifications. Emission from a pulsed 483-nm laser diode (LDH-P-C-470, PicoQuant) was cleaned up (Chroma ET485/20x, F49-482; AHF analysentechnik AG), emission from a 635-nm laser diode (LDH-P-C-635B, PicoQuant) was cleaned up (Chroma z635/10x, PicoQuant), and both lasers were alternated at 26.67 MHz (PDL 828 Sepia II, PicoQuant), delayed ~18 ns with respect to each other, and combined with a 483-nm reflecting dichroic mirror in a single-mode optical fiber (coupler, 60FC-4- RGBV11-47; fiber, PMC-400Si-2.6-NA012-3-APC-150-P, Schäfter + Kirchhoff GmbH). After collimation (60FC-L-4-RGBV11-47, SuK GmbH), the linear polarization was cleaned up (CODIXX VIS-600- BC-W01, F22-601; AHF analysentechnik AG), and the light (75 μW of 483-nm light and 50 μW of 635-nm light) was reflected into the back port of the microscope (IX70, Olympus Belgium NV) and upward [3-mm-thick full-reflective Ag mirror, F21-005 (AHF) mounted in a total internal reflection fluorescence filter cube for BX2/IX2, F91-960; AHF analysentechnik AG] to the objective (UPLSAPO-60XW, Olympus). Sample emission was transmitted through a 3-mm-thick excitation polychroic mirror (Chroma zt470-488/640rpc, F58-PQ08; AHF analysentechnik AG), focused through a 75-mm pinhole (P75S, Thorlabs) with an achromatic lens (AC254-200-A-ML, Thorlabs), collimated again (AC254-50-A-ML, Thorlabs), and spectrally split (Chroma T560lpxr, F48-559; AHF analysentechnik AG). The blue range was filtered (Chroma ET525/50m, F47-525, AHF analysentechnik AG), and polarization was split (PBS251, Thorlabs). The red range was also filtered (Chroma ET705/100m, AHF analysentechnik AG), and polarization was split (PBS252, Thorlabs). Photons were detected on four avalanche photodiodes (PerkinElmer or EG&G SPCM-AQR12/14, or Laser Components COUNT BLUE), which were connected to a time-correlated single-photon counting (TCSPC) device (SPC-630, Becker & Hickl GmbH) over a router (HRT-82, Becker & Hickl) and power supply (DSN 102, PicoQuant). Signals were stored in 12-bit first-in-first-out (FIFO) files. Microscope alignment was carried out using fluorescence correlation spectroscopy (FCS) on freely diffusing ATTO 488-CA and ATTO 655-CA (ATTO-TEC) and by connecting the detectors to a hardware correlator (ALV-5000/EPP) over a power splitter (PSM50/51, PicoQuant) for alignment by real-time FCS. Instrument response functions (IRFs) were recorded one detector at-a-time in a solution of ATTO 488-CA or ATTO 655-CA in near-saturated centrifuged potassium iodide at a 25-kHz average count rate for a total of 25 × 10^6^ photons. Macrotime-dependent microtime shifting was corrected for two (blue/parallel and red/perpendicular) of four avalanche photodiodes (APDs) using the IRF data as input (Otosu et al., 2013).

### Single-molecule MFD-PIE-FRET data analysis

#### Burst Identification

Data was analyzed with the PAM software (Schrimpf et al., 2018) via standard procedures for MFD-PIE smFRET burst analysis (Hellenkamp et al., 2018; Kudryavtsev et al., 2012). Signals from each TCSPC routing channel (corresponding to the individual detectors) were divided in time gates to discern 483-nm excited FRET photons from 635-nm excited acceptor photons. A two-color MFD all-photon burst search algorithm using a 500-μs sliding time window (minimum of 100 photons per burst, minimum of 5 photons per time window) was used to identify single donor- and/or acceptor–labeled molecules in the fluorescence traces. Double labelled single molecules were selected from the raw burst data using a kernel density estimator (ALEX-2CDE < 12) (Tomov et al., 2012), that also excluded other artefacts. Sparse slow-diffusing aggregates were removed from the data by excluding bursts exhibiting a burst duration > 20 ms. By generating histograms of *E* versus measurement time, we corroborated that the distribution of *E* was invariant over the duration of the measurement.

#### FRET efficiency and stoichiometry ratio

The absolute burst-averaged FRET efficiency E_FRET_ was calculated with:

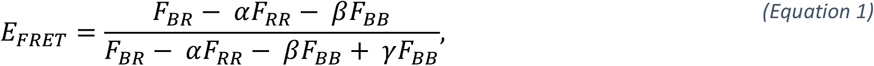

where *F_BR_* = *S_BR_* − *B_BR_* is the background corrected number of photons in both red detection channels after blue excitation. (with *S_BR_* and *B_BR_* the summed intensity and background, respectively, in time gates BR_‖_ and BR_⊥_); *F_BB_* = *S_BB_* – *B_BB_* the background-corrected number of photons in the blue detection channel after blue excitation (with S_BB_ and B_BB_ the summed intensity and background, respectively, in time gates BB_‖_ and BB_⊥_), *F_RR_* = *S_RR_* − *B_RR_* the background corrected number of photons in the red detection channel after red excitation (with *S_RR_* and *B_RR_* the summed intensity and background, respectively, in time gates RR_‖_ and RR_⊥_), α a correction factor for direct excitation of the acceptor with the 483 nm laser, β a correction factor for emission crosstalk of the donor in the acceptor channel, and α the relative detection efficiency of the donor and acceptor. (Kudryavtsev et al., 2012)

The corrected stoichiometry ratio S was calculated with:

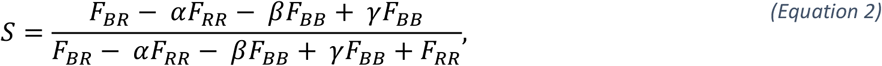

resulting in the ratio of the blue laser excited photons over all excited photons (blue and red laser). According to this calculation D-only labeled molecules will have S values near unity, while A-only labeled molecules will have values near zero. Double labeled molecules will exhibit S values between 0.2-0.6 depending on the used dye pair, the microscope and the laser power ratio.

#### Data correction

First, background was subtracted from the experimental signals. Then, the β- and α-factors were determined directly from the measurement (Kudryavtsev et al., 2012) and data were corrected. Finally, the center values of the *E-S* data cloud for each protein were estimated manually, plotted in an *E* vs. *1/S* graph, and a straight line was fitted to the resulting data to obtain the γ-factor:

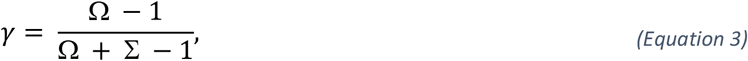

where Ω is the intercept and Σ the slope of the linear fit.

#### Distances

Finally, FRET-averaged D-A distances were obtained from the center *E_FRET_* values with:

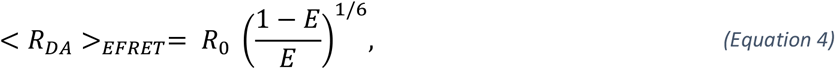

Where *R_0_* is the Förster distance (56.8 Å) (Vandenberk et al., 2018) using the measured dye spectra, a refractive index n = 1.33, an orientation factor κ^2^ = 2/3, a donor quantum yield Φ = 0.6 for ATTO488 as used in (Tamman et al., 2020) and acceptor extinction coefficient ε = 265,000 cm^−1^M^−1^. The quantum yield was determined using a home-built absorbance/fluorescence spectroscope (Moeyaert et al., 2014). For simplicity, *R_DA_* will be noted *R* throughout the text.

#### Burstwise fluorescence lifetimes

A maximum likelihood estimator approach (MLE (Schaffer et al., 1999)) was used to estimate single-molecule burst-averaged single exponential fluorescence lifetimes of the FRET donor in presence of the acceptor, *τ_D (A)_*, and FRET acceptor, *τ_A_*. For molecules that are conformationally static during transit through the laser focus, the FRET efficiency is related to the fluorescence lifetime of the donor as follows:

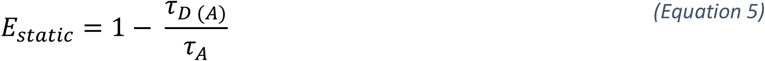

However, dyes are attached to the molecule of interest via flexible linkers, resulting in a Gaussian D-A distance distribution, even for conformationally static molecules. Especially at short distances (high FRET), this effect causes a non-linear relation between the intensity-based *E_FRET_* and *τ_D (A)_*. We simulated this ‘static FRET line including linker dynamics’ as follows: we calculated *m* values for *R* between 0 and 3 x *R_0_*. For every R, we calculated a Gaussian distribution of *p* distances around the central *R*, with the apparent linker length as the standard deviation, resulting in a list of *m*p* values for *R*. For every *R*, we calculated which donor fluorescence lifetime would be associated with it ((Equation 5), with *τ_D_* the mean burstwise lifetime of raw burst data with *S* > 0.8). The apparent linker length (5.6 Å) was obtained from a sub-ensemble donor fluorescence lifetime fitting of double-labeled molecules using a Gaussian distance distribution model. Finally, we calculated the species-weighted average lifetime, and from it the intensity-based *E_FRET_* (x-axis of the static FRET line) and the intensity weighted average lifetime (y-axis of the static FRET line).

Similarly, for molecules exhibiting multiple lifetimes during transit due to conformational FRET dynamics, the burst-averaged lifetime will be fluorescence-weighted towards the long-lifetime species that emits more photons, resulting in an even further upward shift of the experimental data from the theoretical line (*Equation 5*).

### Photon distribution analysis (PDA)

Static PDA was carried out to obtain the absolute interdye distance distribution as described before, assuming two Gaussian distributed states (Antonik et al., 2006). Practically, for each FRET data set, raw bursts were re-binned in 1 ms time bins, and histograms were constructed and analyzed. Binned data were plotted in a raw (uncorrected) FRET efficiency (*E*_PR_) versus uncorrected stoichiometry (*S*_PR_) plot and only the bins with 0.2 < *S*_PR_ < 0.6 were included in the analysis to remove burst sections containing complex acceptor photophysics or photobleaching. Furthermore, only bins with at least 20 and maximally 200 photons (to reduce calculation time) were used for PDA analysis. A two-state model for a Gaussian distance distribution was used to generate a library of simulated *E*_FRET_ values, which was subsequently fitted to the experimental *E*_FRET_ histogram using a reduced χ^2^–guided simplex search algorithm to obtain the amplitude, mean distance *R* and width s of all Gaussian distributed substates and, in the case of multiple states, their area fraction *A* (%). A probability density function (PDF) was calculated per state using the R and σ parameters obtained from PDA analysis that describe the underlying Gaussian distributed states. The summed PDF was scaled to a total area of unity, with the PDF area of each state scaled to the corresponding fraction of molecules. Criteria for a good fit were a low reduced χ2 value, as well as a weighted-residual plot free of trends.

## Acknowledgements

We thank C.R. O’Riordan for C127 cells and J.R. Riordan, T. Jensen and CF Foundation Therapeutics for anti-CFTR antibodies. We thank John Kappes for HEK293 CFTR cells. C.G. acknowledges support by the Fond Forton, the Welbio (grant CR-2012S-04R), Vaincre la Mucoviscidose, Mukoviszidose e.V., the Association Luxembourgeoise de Lutte contre la Mucoviscidose, ABCF2, the Chiesi Fondation, the Cystic Fibrosis Foundation, the Fondation Air Liquide and the Fondation ULB. D.N.S acknowledges support from the CF Trust and CF Foundation Therapeutics.

We are grateful to H. Remaut for careful reading of the manuscript. We acknowledge Diamond Light Source for time on Beamlines i02, i04 and i24 under Proposals 12718 and 9426.

J.S. acknowledges support from Instruct-ERIC, part of the European Strategy Forum on Research Infrastructures (ESFRI), Instruct-ULTRA (EU H2020 Grant 731005), and the Research Foundation - Flanders (FWO) for support with nanobody discovery. C.G. is a senior Research Associate of the FRS-FNRS. D.S. was a fellow of the FRIA.

We thank Frank Sobott and James Ault of the Biomolecular Mass Spectrometry department in the Astbury Centre for their support and assistance in this work, and the BBSRC (BB/M012573/1) for funding.

## Author contributions

D.S., D.N.S. and C.G. conceived and designed the experiments. D.S., M.S., M.O., Y.W., R.C., E.P., T.L. and C.M. performed the research. D.S, M.S., M.O., Y.W., C.M., A.G.P., D.N.S., J.H., and C.G. analysed the data. D.S., D.N.S., J.S., J.H. and C.G. co-wrote the manuscript. D.N.S., J.S., J.H., and C.G. directed the study. All authors approved the final version of the manuscript.

## Data availability

Data supporting the findings of this manuscript are available from the corresponding author upon reasonable request. The atomic coordinates and structure factors reported in this paper have been deposited in the Protein Data Bank (PDB). The accession numbers for the structure reported in this paper is PDB: 6ZE1.

