## Supplementary Material for "A Topological Switch Enables Misfolding of the Cystic Fibrosis Transmembrane Conductance Regulator"

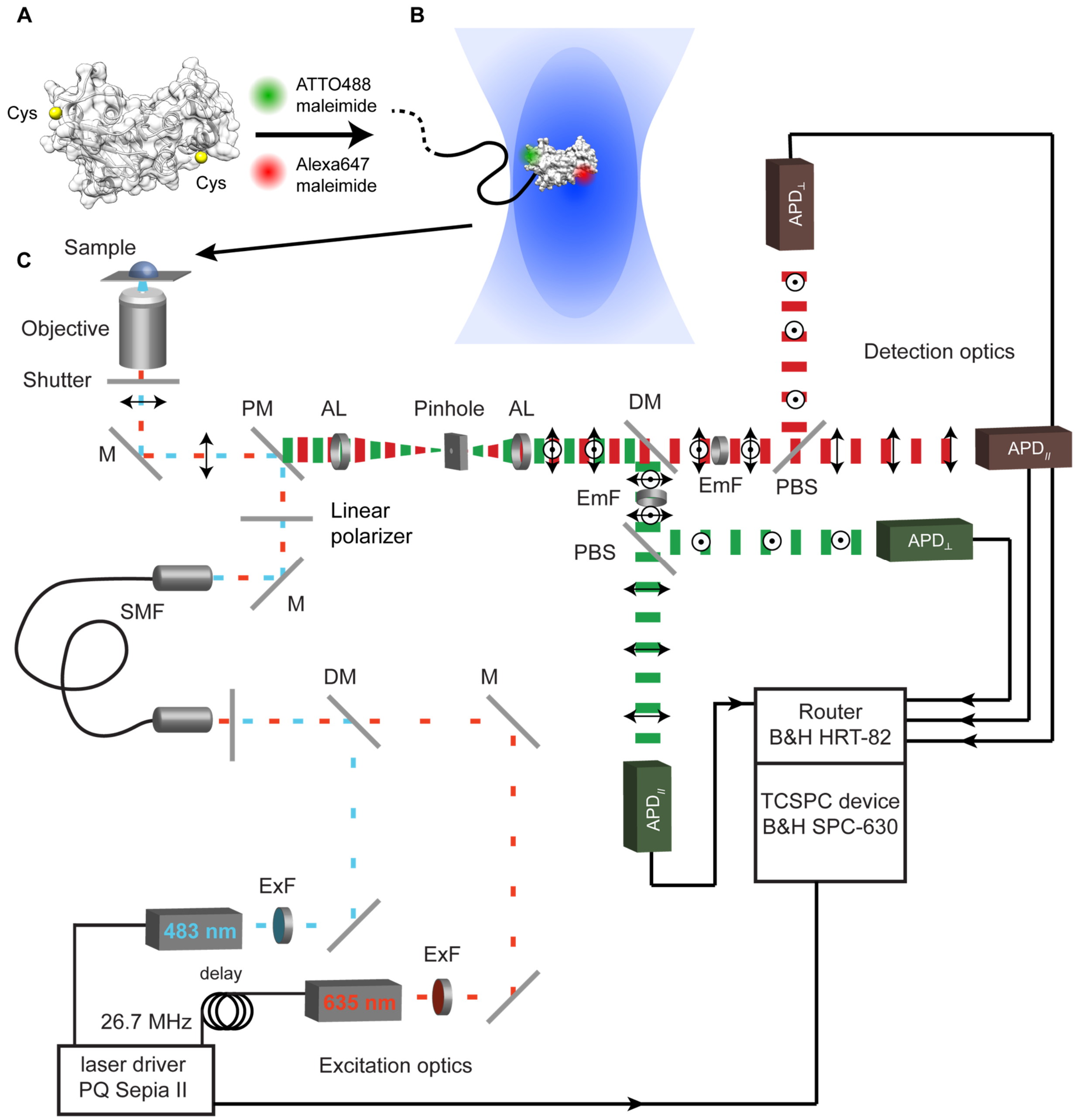


**Fig. S1 | Setup of the two-color MFD-PIE microscope.**

**A,** Representation of NBD1 containing two cysteines residues (yellow spheres) which are labeled by the fluorophores ATTO488 and Alexa647. **B,** FRET between both fluorophores attached to NBD1 is monitored while NBD1 diffuses through the confocal volume. **C,** Schematic illustration of the confocal microscope setup used here to study molecules freely diffusing in solution by multiparameter fluorescence detection with pulsed interleaved excitation (MFD-PIE) (Hellenkamp et al., 2018; Kudryavtsev et al., 2012). The lasers emit pulses with a fixed repetition rate and are synchronized by the TSCPS hardware (see below). An electronic delay leads to a lag time between green and red laser leading to interleaved excitation of the sample. Dichroic mirrors (DM) combine the lasers. They are then collimated and their beam profile is cleaned up by a single-mode optical fiber (SMF). Polychroic mirrors (PM) separate the excitation from the emission light both spectrally and spatially. The emission light is focused through a confocal pinhole and collimated by achromatic lenses (AL) to reduce chromatic aberrations. The emission light is then split up into two spectral ranges (green and red). Polarization beam splitters (PBS) further split the polarization of each color range into parallel (‖) and antiparallel (⊥). Emission filters (EmF) finally transmit the appropriate spectral band and a lens focuses the emission light onto an avalanche photodiode (APD). The time-correlated single photon counting (TCSPC (Becker et al., 1999)) hardware receives the signals from the APDs. Three parameters are registered per photon: the identity of the detector (red or green, parallel or antiparallel), the microtime (time since the last laser pulse) and the macrotime (time since the beginning of the experiment).

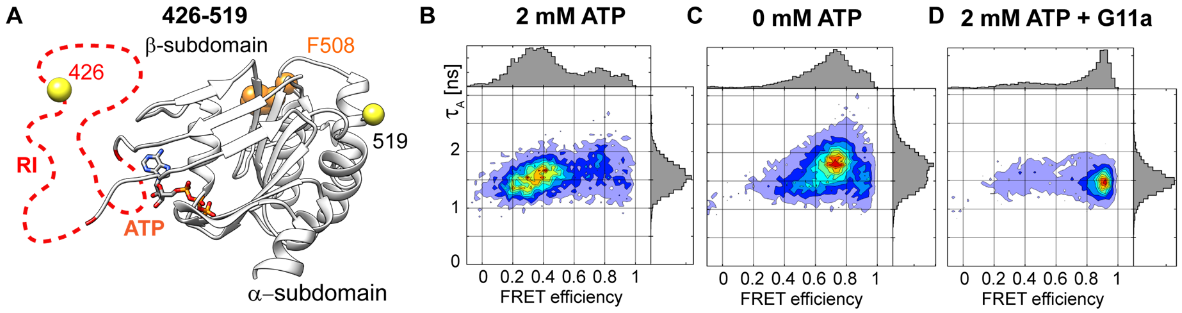


**Fig. S2 | Acceptor fluorescence lifetime versus FRET efficiency plots for the 426-519 reporter pair.**

**A,** Location of the smFRET reporters in the canonical conformation of NBD1 (based on the published structure of human NBD1, PDB: 2BBO). The positions of the labelled cysteines are shown as yellow spheres. The regulatory insertion (RI), which is not resolved in the structure, is depicted as a red dashed line. F508, located in the α-subdomain, and ATP are shown. **B-D,** Two-dimensional histograms of acceptor fluorescence lifetime ($\tau_A$) versus FRET efficiency of the 426-519 reporter for the indicated conditions. The histograms provide a distinctive signature for each of the three observed FRET states. Small changes in acceptor lifetimes are due to an altered fluorescence quantum yield of the acceptor, likely due to sticking of the fluorophore to the protein. Correction of these effects did not substantially affect the observed FRET efficiency values (data not shown).

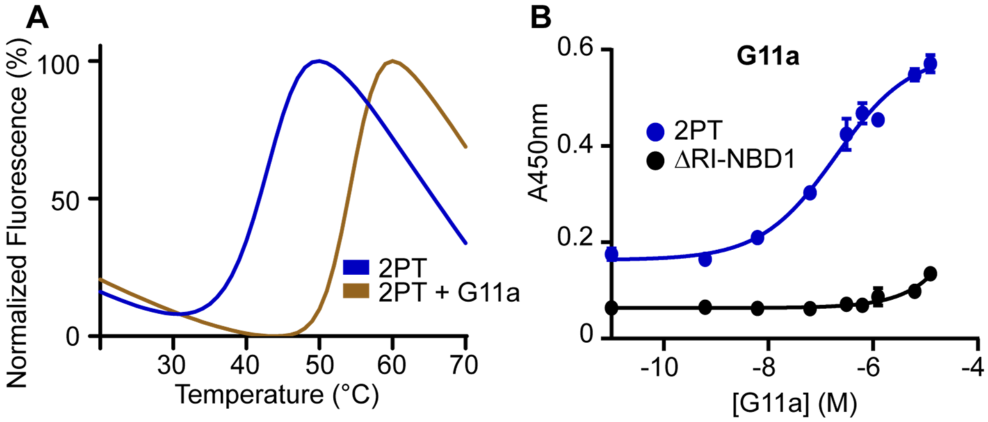


**Fig. S3 | Characterization of the interactions between NBD1 variants and nanobody G11a.**

**A,** Differential scanning fluorescence (DSF) of purified NBD1 variants with and without nanobody G11a. The samples were incubated with SYPRO Orange dye and fluorescence was measured as a function of temperature. The melting temperatures ($T_m$) were determined by the maxima of the first derivative of fluorescence. Curves depict mean of duplicates of one experiment representative of at least three independent experiments. **B**, Binding of nanobody G11a to NBD1 variants measured by ELISA. Biotinylated NBD1 variants were immobilized on neutravidin-coated plates and incubated with increasing concentrations of nanobody. Binding of nanobody was followed by immunodetection of the His6-tag (see Methods). Representative curve of 3 independent experiments is shown. Error bars represent the standard deviation (SD) of duplicates. See Table S1 for $pEC_{50}$ and $T_m$ values.

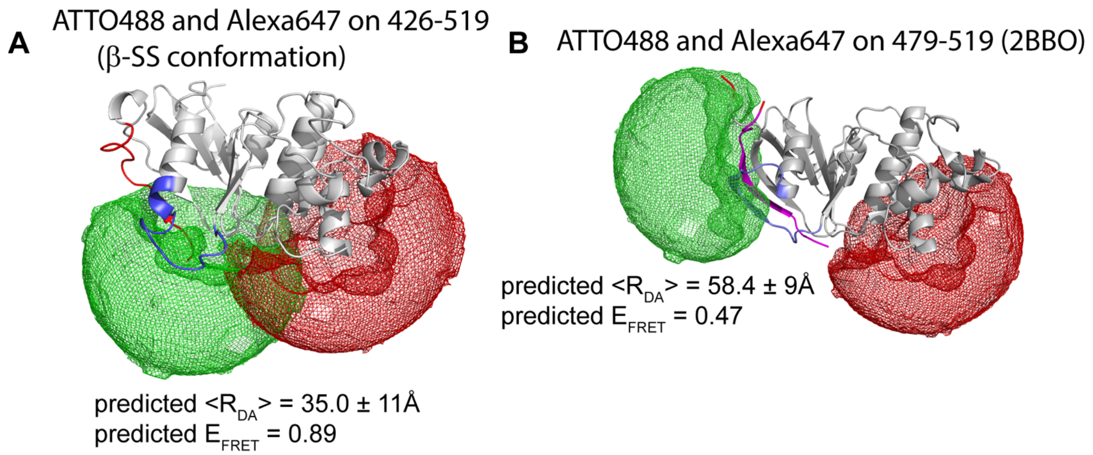


**Fig. S4 | FRET-restrained positioning and screening simulation.**

FRET-restrained positioning and screening (FPS (Kalinin et al., 2012)) simulations of the accessible volumes of the donor and acceptor dyes ATTO488 and Alexa647 when attached to cysteine residues in NBD1. **A**, Accessible dye positions when attached to residues 426 and 519 in the β-strand swapped conformation observed in the 2PT:G11a structure. **B**, Accessible dye positions in the 479-519 reporter in the canonical conformation, based on the canonical NDB1 structure (PDB: 2BBO). Simulation results for all used reporters in both β-strand swapped and canonical conformations are detailed in Table S3). Figures were generated using PyMOL (DeLano Scientific LLC).

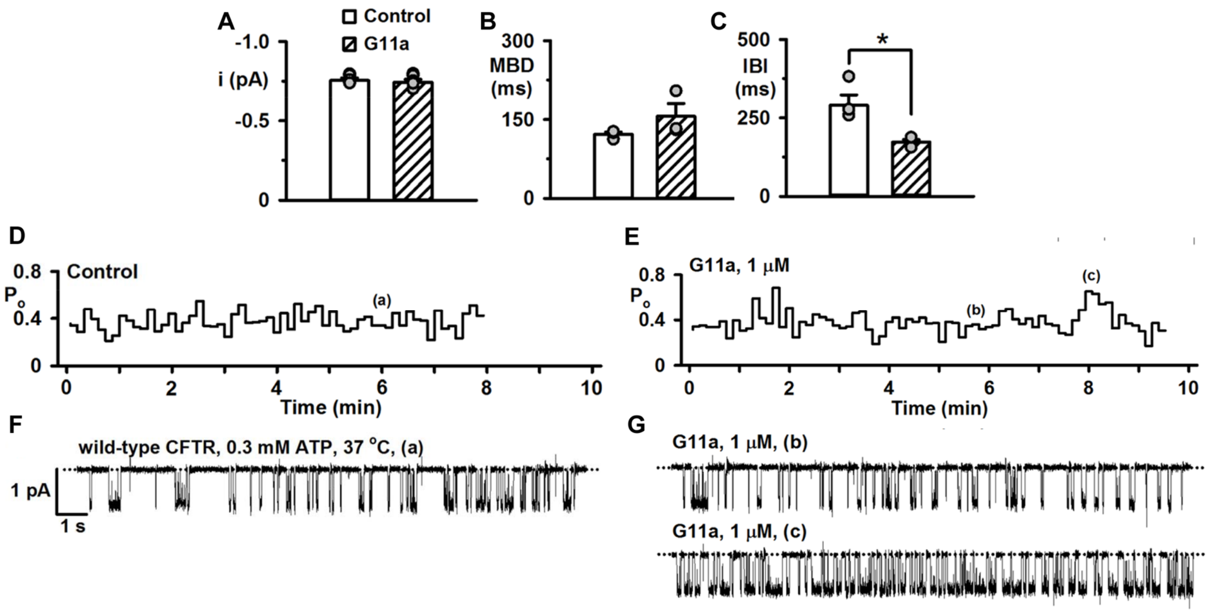


**Fig. S5 | G11a binds wild-type human CFTR and alters its gating behaviour.**

**A-C,** Summary of patch-clamp studies of wild-type CFTR $Cl^-$ channel in an excised inside-out membrane patch from a C127 cell heterologously expressing wild-type human CFTR. The data were acquired at 37 °C in the presence of ATP (0.3 mM) and PKA (75 nM) in the intracellular solution. After the channel was fully activated, G11a (1 µM) was then directly added to the intracellular solution bathing the membrane patch. A large $Cl^-$ concentration gradient was imposed across the membrane patch ($[Cl^-]_{int}$, 147 mM; $[Cl^-]_{ext}$, 10 mM) and membrane voltage was clamped at –50 mV. Single-channel current amplitude (i) mean burst duration (MBD) and interburst interval (IBI) of wild-type CFTR in the absence and presence of G11a are quantified. Symbols represent individual values and columns are means ± SEM (n = 6); *, P < 0.05 vs control. **D-E**, Open probability ($P_o$) timecourses for an individual wild-type CFTR $Cl^-$ channel in absence and presence of G11a (1 µM). Lowercase letters indicate the locations of the recordings shown in **F** and **G** during the $P_o$ timecourses. **F-G**, Representative recordings of an individual wild-type CFTR $Cl^-$ channel in the absence and presence of G11a. The two recordings in the presence of G11a demonstrate the different gating patterns observed in its presence (b, steady-state gating; c, high activity gating). Dotted lines indicate where the channel is closed and downward deflections correspond to channel openings.

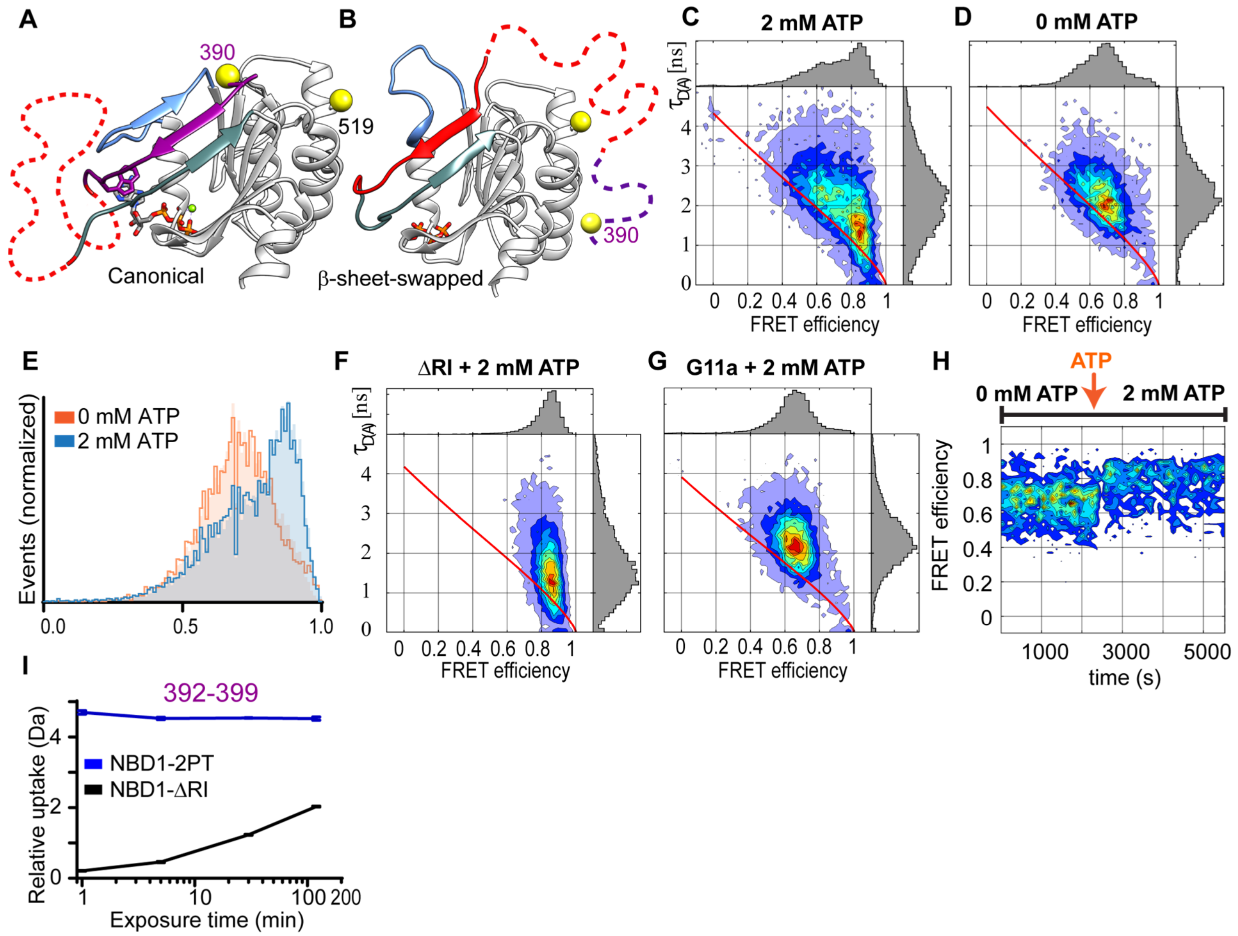


**Fig. S6 |** Conformational dynamics of the 390-519 reporter pair.

**A-B.** Structures of the canonical conformation (PDB: 2BBO) and the β-strand swapped conformation reported here. Unresolved segments are depicted in dashed lines, the RI (404-436) in red and the N-terminal region (389-403) in magenta. **C-D,** Donor lifetime in presence of acceptor ($\tau_{D\,(A)}$) vs FRET efficiency histograms of the 390-519 reporter pair with and without ATP. The red line is called the 'static FRET line' and characterizes the theoretical relationship between lifetime and FRET efficiency in absence of conformational dynamics on the timescale of the measurement (milliseconds). **E**, Overlay of FRET efficiency histograms of 2PT-390-519 without ATP and with 2 mM ATP. Raw data (fill) was fitted using the PDAFit software (thick lines, see Methods). **F-G,** Donor lifetime vs FRET efficiency plots of 2PT-390-519 without the RI and with the RI in presence of nanobody G11a. **H**, Reversibility of the conformational change was tested. At the beginning of the experiment no ATP was present and after 2500s ATP was added to yield a final concentration of 2 mM. **I,** Deuterium build-up curves of the 392-399 peptide identified in mass spectral analyses of NBD1-2PT (blue) and ΔRI-NBD1 (black).

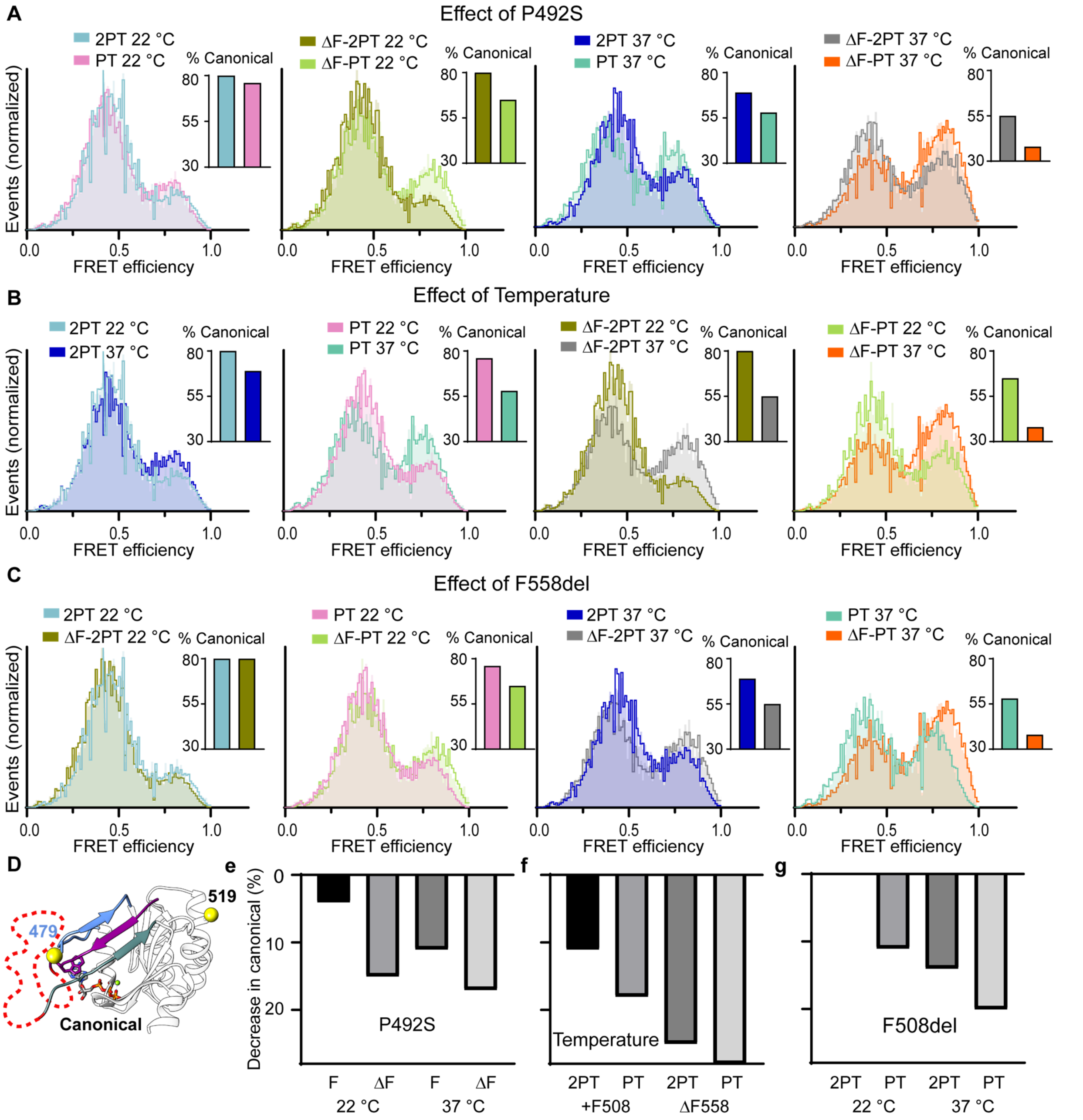


**Fig. S7** | Effects of the P492S substitution, temperature and F508del on the conformational equilibrium measured via the 479-519 reporter pair.

**A-C**, Overlay of FRET efficiency histograms of different 479-519 reporter pair variants. Raw data (fill) was fitted using the PDAFit software (thick lines, see Methods). The insets show the population of the canonical state. **D**, Illustration of the distance reporter pair 479-519. **E-G**, Decrease in the population of the canonical state upon substitution of P492 by the wild-type Serine (**E**), upon increasing the temperature from 22 °C to 37 °C (**F**) and upon deletion of F508 (**G**).

**Table S1 | $EC_{50}$, $T_m$ and $\Delta T_m$ values for nanobody G11a in complex with NBD1 variants**

| | $pEC_{50}$ (M) | $T_m$ (°C) | $\Delta T_m$ (°C) |
|---|---|---|---|
| 2PT | / | 43.8 ± 0.3 | / |
| G11a + 2PT | 6.9 ± 0.1 | 53.8 ± 0.6 | + 10 |
| G11a + ΔRI | N.A. | / | / |

pEC50 and Tm values are mean ± SEM from at least three independent experiments.

**Table S2 | Data collection and processing.**

The $CC_{1/2}$ criterion was used to determine the resolution range. Values for the outer shell are given in parentheses.

| **Sample** | **hNBD1-G11a** |
|---|---|
| Diffraction source | Soleil PX1 |
| Wavelength (Å) | 0.9786 |
| Space group | $P6_122$ |
| *a*, *b*, *c* (Å) | 126.9 126.9 117.4 |
| α, β, γ (°) | 90.0 90.0 120.0 |
| Resolution range (Å) | 109.94 - 2.70 (3.08 - 2.70) |
| Total N°. of reflections | 97389 (3618) |
| N°. of unique reflections | 7989 (399) |
| Completeness ellipsoidal (%) | 93.3 (80.0) |
| Redundancy | 12.2 (9.1) |
| $\langle I/\sigma(I) \rangle$ | 8.0 (1.9) |
| $CC_{1/2}$ | 0.996 (0.647) |
| $R_{r.i.m.}$ | 0.297 (1.425) |
| $R_{pim}$ | 0.09 (0.48) |
| *B* factor (Wilson plot ($Å^2$)) | 73.9 |
| R-factor (%) | 20.7 |
| $R_{free}$-factor (%) | 25.8 |
| Ramachandran profile | |
| Core | 94.0 |
| Allowed | 6.0 |
| Outliers | 0.0 |
| R.m.s. deviations | |
| Bond lengths (Å) | 0.008 |
| Bond angles (°) | 0.98 |
| Number of atoms | 2558 |
| Macromolecules | 2485 |
| Solvent | 41 |
| Ligands | 32 |
| B-factors (Å2) | 71.1 |
| PDB ID | 6ZE1 |

**Table S3 | Modeling of dye distributions using the FPS simulation program**

| CANONICAL (based on 2BBO) | | | | |
|---|---|---|---|---|
| Reporter | $R_{mp}$ (Å) | $\sigma_{DA}$ (Å) | $E_{FRET}$ | $\langle R_{DA} \rangle_{EFRET}$ |
| 390-519 | 29.6 | 10.5 | 0.923 | 37.6 |
| 426-519 | / | / | / | / |
| 442-519 | 55.8 | 8.6 | 0.477 | 57.7 |
| 479-519 | 55.8 | 9 | 0.474 | 57.8 |
| β-STRAND SWAPPED (based on G11a:2PT structure) | | | | |
| Reporter | $R_{mp}$ (Å) | $\sigma_{DA}$ (Å) | $E_{FRET}$ | $\langle R_{DA} \rangle_{EFRET}$ |
| 390-519 | / | / | / | / |
| 426-519 | 31.1 | 11.4 | 0.890 | 40.1 |
| 442-519 | 56.5 | 7.9 | 0.458 | 58.4 |
| 479-519 | 47.5 | 9.3 | 0.644 | 51.5 |

$R_{mp}$ designates the distance between mean dye positions and $\langle RDA \rangle_{EFRET}$ designates the distance formally calculated from simulated FRET efficiencies. For more details on the FPS simulation program see methods.

**Table S4 | Photon distribution analysis (PDA) results of the 479-519 reporter**

| 479-519 | Can | $R_{can}$ (Å) | $\sigma_{can}$ (Å) | Alt | $R_{alt}$ (Å) | $\sigma_{alt}$ (Å) | $\chi$2 |
|---|---|---|---|---|---|---|---|
| 2PT-479-519 + 2 mM ATP 22 °C | 0.80 | 59.5 | 2.3 | 0.20 | 45.9 | 3.7 | 5.8 |
| 2PT-479-519 + 2 mM ATP + G11a 22 °C | 0.19 | 61.2 | 5.2 | 0.81 | 46.7 | 3.7 | 12.52 |
| 2PT-479-519 + 0.2 mM ATP 22 °C | 0.58 | 60.2 | 3.6 | 0.42 | 46.0 | 2.6 | 3.0 |
| 2PT-479-519 without ATP 22 °C | 0.25 | 58.0 | 2.9 | 0.75 | 44.8 | 2.9 | 2.9 |
| dRI-2PT-479-519 + 2mM ATP 22 °C | 1 | 58.7 | 2.9 | | | | 12.3 |
| F508del-2PT-479-519 + 2mM ATP 22 °C | 0.80 | 60.5 | 2.3 | 0.2 | 47.7 | 4.4 | 5.6 |
| F508del-PT-479-519 + 2 mM ATP 22 °C | 0.65 | 60.4 | 2.3 | 0.35 | 46.1 | 4.1 | 2.4 |
| PT-479-519 + 2 mM ATP 22 °C | 0.76 | 60.4 | 2.5 | 0.24 | 47.0 | 3.2 | 4.1 |
| 2PT-479-519 + 2 mM ATP 37 °C | 0.69 | 59.9 | 2.1 | 0.31 | 46.9 | 3.7 | 4.0 |
| PT-479-519 + 2 mM ATP 37 °C | 0.58 | 62.2 | 2.0 | 0.42 | 48.2 | 3.3 | 6.5 |
| F508del-2PT + 2 mM ATP 37 °C | 0.55 | 61.6 | 1.9 | 0.45 | 46.7 | 4.6 | 2.4 |
| F508del-PT-479-519 + 2 mM ATP 37 °C | 0.38 | 61.0 | 2.3 | 0.62 | 46.1 | 4.9 | 4.4 |

Photons from each burst were used to generate shot-noise limited FRET efficiency histograms fitted using the Histogram Library fit method (see Methods). Fraction of molecules in canonical conformation, corresponding to the low FRET state, or in alternative conformations, corresponding to the high FRET state, are given by 'Can' and 'Alt', respectively. Distance and standard deviation of each population are given by R and $\sigma$.
